# A liquid-like coat mediates chromosome clustering during mitotic exit

**DOI:** 10.1101/2023.11.23.568026

**Authors:** Alberto Hernandez-Armendariz, Valerio Sorichetti, Yuki Hayashi, Andreas Brunner, Jan Ellenberg, Anđela Šarić, Sara Cuylen-Haering

## Abstract

The individualization of chromosomes during early mitosis and their clustering upon exit from cell division are two key transitions that ensure efficient segregation of eukaryotic chromosomes. Both processes are regulated by the surfactant-like protein Ki-67, but how Ki-67 achieves these diametric functions has remained unknown. Here, we report that Ki-67 radically switches from a chromosome repellent to a chromosome attractant during anaphase. We show that Ki-67 dephosphorylation during mitotic exit and the simultaneous exposure of a conserved basic patch induce the RNA-dependent formation of a liquid-like condensed phase on the chromosome surface. Experiments and coarse-grained simulations support a model in which the coalescence of chromosome surfaces driven by phase separation promote clustering of chromosomes. Our study reveals how the switch of Ki-67 from a surfactant to a liquid-like condensed phase can generate the mechanical forces during genome segregation that are required for re-establishing nuclear-cytoplasmic compartmentalization after mitosis.

## INTRODUCTION

Cell division requires a dramatic reorganization of the genome. During entry into mitosis, the formerly loose network of chromatin fibers compacts into dense mitotic chromosomes, built from consecutive chromatin loops that are held together by proteins in a network-like manner^1^. Nuclear envelope breakdown releases mitotic chromosomes into the cytoplasm^2^ and their surface becomes covered by an intricate layer of proteins and ribonucleic acids (RNAs), many of which are otherwise found in the nucleolus during interphase^3–5^. Recent studies have demonstrated that the proliferation marker protein Ki-67 is central to the organization of the chromosome surface, since the entire surface layer is delocalized in its absence^6–8^.

During early mitosis, Ki-67 relocalizes from nucleoli to the chromosome surface^9^, where it forms extended brush structures ∼90 nm in length^8^. These brushes are thought to mediate the dispersion of individual mitotic chromosomes for their attachment to mitotic spindle microtubules^8^. Individualized chromosomes are then captured by mitotic spindle microtubules and, once all chromosomes have been bioriented, anaphase segregation is triggered by cleavage of cohesin^10^. During exit from mitosis, the Ki-67 brush structure collapses, and chromosomes cluster in a Ki-67-dependent manner to facilitate exclusion of large cytoplasmic particles from the future nuclear space^11^. These findings suggest that Ki-67 must fundamentally switch its function between mitotic entry and exit, but the underlying mechanistic basis has remained elusive.

Ki-67 contains several features that have been linked to liquid-liquid phase separation, including structural disorder, low-complexity regions, and multivalency^12^. A recent study^13^ reported that single or multiple copies of recombinantly expressed Ki-67 repeats can undergo phase separation *in vitro* at high protein concentrations and in the presence of crowding agents. Phase separation was notably amplified upon phosphorylation and it has been proposed in this study that phosphorylation plays a pivotal role in Ki-67 phase separation by generating segments with alternating charges within the repeat modules, referred to as “charge blockiness”. Since Ki-67 is phosphorylated upon entry into mitosis^14^, it implies that Ki-67 forms a liquid-like layer on the surface of chromosome during early mitosis. However, direct evidence supporting this hypothesis in living cells is missing, as are insights into the possible significance and relevance of Ki-67 liquid-like properties or its dependency on the cell cycle stage.

Here, we combine quantitative live-cell imaging and protein engineering experiments with data-driven coarse-grained simulations to investigate the change of Ki-67’s properties through the course of mitosis to understand how chromosomes can completely invert their physical properties. We demonstrate that, upon dephosphorylation at anaphase onset, Ki-67 phase separates into a liquid-like phase and that this transition is RNA-dependent. Consequently, chromosome surfaces no longer repulse but instead attract each other, thereby promoting chromosome clustering and the subsequent exclusion of cytoplasm from the newly formed nucleus. Our findings hence illustrate how phase separation can generate large-scale adhesive forces to perform essential cellular functions.

## RESULTS

### Ki-67 changes its properties during mitotic exit

To investigate properties of Ki-67 during mitosis and unravel their role in chromosome clustering, we utilized a previously established assay that triggers mitotic exit in the absence of a spindle but nevertheless facilitates timely assembly of a sealed and transport-competent nuclear envelope^11^. This assay is based on first inducing mitotic arrest using nocodazole, which depolymerizes microtubules, followed by acute inhibition of the mitotic kinase CDK1 using flavopiridol^15^.

As expected, chromosomes clustered within a few minutes after addition of flavopiridol. The total signal intensity of chromosome-bound Ki-67 increased concomitantly (Figures 1A and 1B; Video S1). This increase was also obvious in a cell line that expresses endogenously tagged Ki-67 (Figures S1A and S1B). Furthermore, FCS-calibrated imaging of unperturbed cells revealed that Ki-67 endogenous concentration on chromosomes increased from ∼500 nM in metaphase to 750 nM in anaphase (Figures S1D–S1I). These observations contradict the possibility that chromosome clustering is simply triggered by a reduction of the chromosomal fraction of Ki-67.

**Figure 1.**
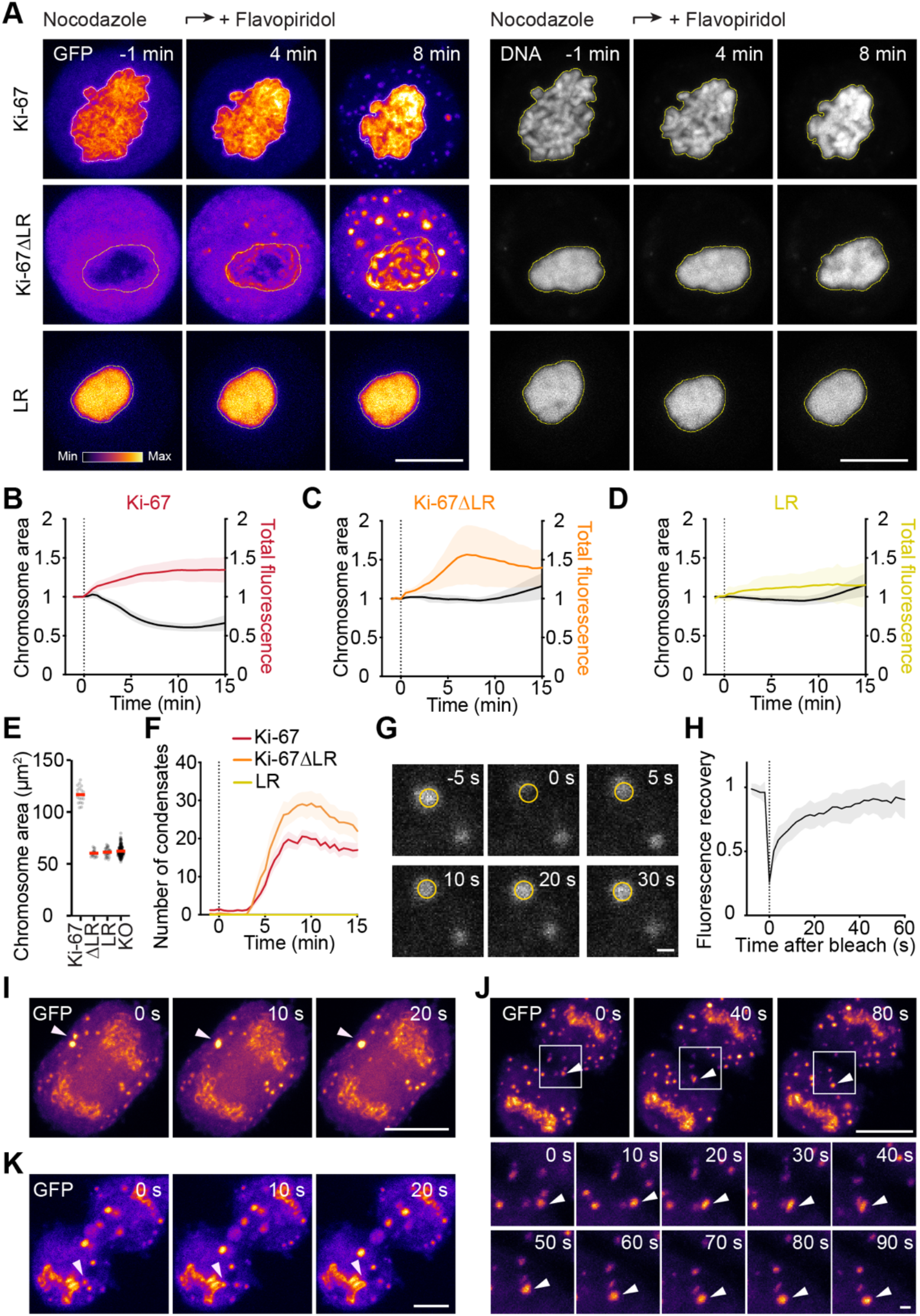
Ki-67 phase separates and enriches on chromosomes as they cluster during mitotic exit. (A) Time-lapse microscopy images of spindle-less mitotic exit of HeLa Ki-67 KO cells transiently expressing Ki-67 full-length, Ki-67ΔLR or Ki-67’s LR domain tagged with EGFP. Cells were imaged in the presence of nocodazole; flavopiridol was added (t = 0 min) to induce mitotic exit. DNA was labelled with SiR-Hoechst. White and yellow lines represent chromosomal segmentations quantified in (B-D). (B–D) Quantification of chromosome ensemble area over time and EGFP total fluorescence intensity in the chromosomal area as in (A). (E) Quantification of chromosome ensemble area (prior to flavopiridol addition) of Ki-67 KO cells transfected with either EGFP-Ki-67, EGFP-Ki-67ΔLR or EGFP-LR plotted in (B-E). Untransfected Ki-67 KO cells arrested by nocodazole in early mitosis are plotted for reference. n = 21 (Ki-67); n = 18 (Ki-67ΔLR); n = 31 (LR); n = 177 (KO). (F) Quantification of number of cytoplasmic condensates over time as in (A). (G) Individual condensates in Ki-67 KO cells expressing EGFP-Ki-67ΔLR were photobleached and the recovery of fluorescence was followed by time-lapse recording in a circular image region (yellow circle). Representative example of the quantification in (H). (H) Quantification of EGFP mean fluorescence intensity in the bleached area, normalized to pre-bleached values. n = 13. (I) Time-lapse microscopy images of Ki-67 wild-type cell overexpressing EGFP-Ki-67ΔLR undergoing anaphase. Two condensates fusing with each other are indicated by white arrows. (J) Ki-67 KO cell transiently expressing EGFP-Ki-67ΔLR undergoing cytokinesis. Zoom-in images (bottom). (K) Ki-67 wild-type cell overexpressing Ki-67ΔLR-mNeonGreen. White arrows indicate a condensate fusing with the chromosome surface pool. Dashed vertical lines indicate flavopiridol addition (B–D, F) or photobleaching (H). Red bars indicate mean (E). Lines and shaded areas represent mean ± SD (B–D, H) or mean ± SEM (F). Maximum intensity z-projections (A, I, J) or single z-slices (G, K) are shown. Scale bars, 10 µm (A, I, J top, K), 1 µm (G, J bottom).

Remarkably, the chromosomal levels of a Ki-67 version that lacks the DNA-binding LR domain (Ki-67ΔLR)^16^ and therefore localizes to the cytoplasm during early mitosis similarly increased upon mitotic exit (Figures 1A and 1C; Video S1). In contrast, the chromosomal levels of only the LR domain (LR) did not increase upon mitotic exit (Figures 1A and 1D; Video S1). The observed chromosomal recruitment of Ki-67ΔLR cannot be due to its multimerization with endogenous full-length Ki-67, since the constructs were expressed in Ki-67 knock-out (KO) cells. Furthermore, Ki-67 increase on chromosomes preceded binding of the DNA-bridging barrier-to-autointegration factor (BAF)^17^ and the reformation of the nuclear envelope (Figures S1J–S1L), ruling out that it is a consequence of either of these two events.

Neither the Ki-67ΔLR nor the LR construct restored chromosome dispersal during early mitosis in Ki-67 KO cells^8^ (Figures 1A and 1E). Thus, chromosomes remained in a clustered state with a constant area until their decompaction started approximately 10 min after induction of mitotic exit. However, the enrichment of the Ki-67ΔLR construct on chromosomes correlated with chromosome clustering (compare Figures 1B and 1C). Our data therefore suggest that Ki-67 gains a new DNA-binding activity that is independent of the known LR domain during mitotic exit, which might be responsible for triggering chromosome clustering by connecting neighboring chromosomes.

Concomitant with Ki-67ΔLR enrichment on mitotic chromosomes, the soluble pool of Ki-67ΔLR formed micron-sized spherical foci (Figures 1A and 1F; Video S1). Full-length Ki-67 formed foci with similar kinetics, albeit lower in number and size, irrespective of whether it was overexpressed (Figures 1A and 1F) or expressed at endogenous levels (Figures S1A, S1C, S1G, and S1I). The number of foci reached peak levels by the time chromosomes were maximally clustered.

Ki-67ΔLR molecules within foci were highly dynamic, since their fluorescence recovered within 30 seconds after photobleaching (Figures 1G and 1H), and since foci readily fused and relaxed into larger spherical structures over time (Figure 1I). Furthermore, Ki-67ΔLR foci located in the cleavage furrow during cytokinesis deformed due to mechanical stress and rounded up again after relaxation (Figure 1J). In addition, cytoplasmic Ki-67ΔLR foci not only fused with each other (Figure 1I) but also with the chromosome-bound pool of Ki-67ΔLR (Figure 1K; Video S2). These findings and the fact that full-length Ki-67 molecules on the chromosome surface are highly mobile during mitotic exit (Figures S1M and S1N; ^18^) suggest that both the chromosome surface and the cytoplasmic pools of Ki-67 gain liquid-like properties during mitotic exit. Chromosome clustering could thus be driven by a phase transition of Ki-67 into a liquid-like protein layer, by chromosome bridging through the formation of a second DNA-binding site, or by a combination of both.

### A phospho-switch controls chromosome clustering

Although Ki-67 gets highly phosphorylated during early mitosis^14^ at more than one hundred identified sites^19^, its phosphorylation is largely reverted by the end of anaphase^18,20^, possibly through recruitment of protein phosphatase 1 (PP1)^6^. To test whether dephosphorylation triggers the increase of Ki-67 on chromosomes and the change in its phase-separation properties, we generated a phosphomimetic version of Ki-67 by substituting 139 of the serine or threonine residues that had been reported to be phosphorylated^19^ for glutamate residues (Ki-67(139E); Figure 2A).

**Figure 2.**
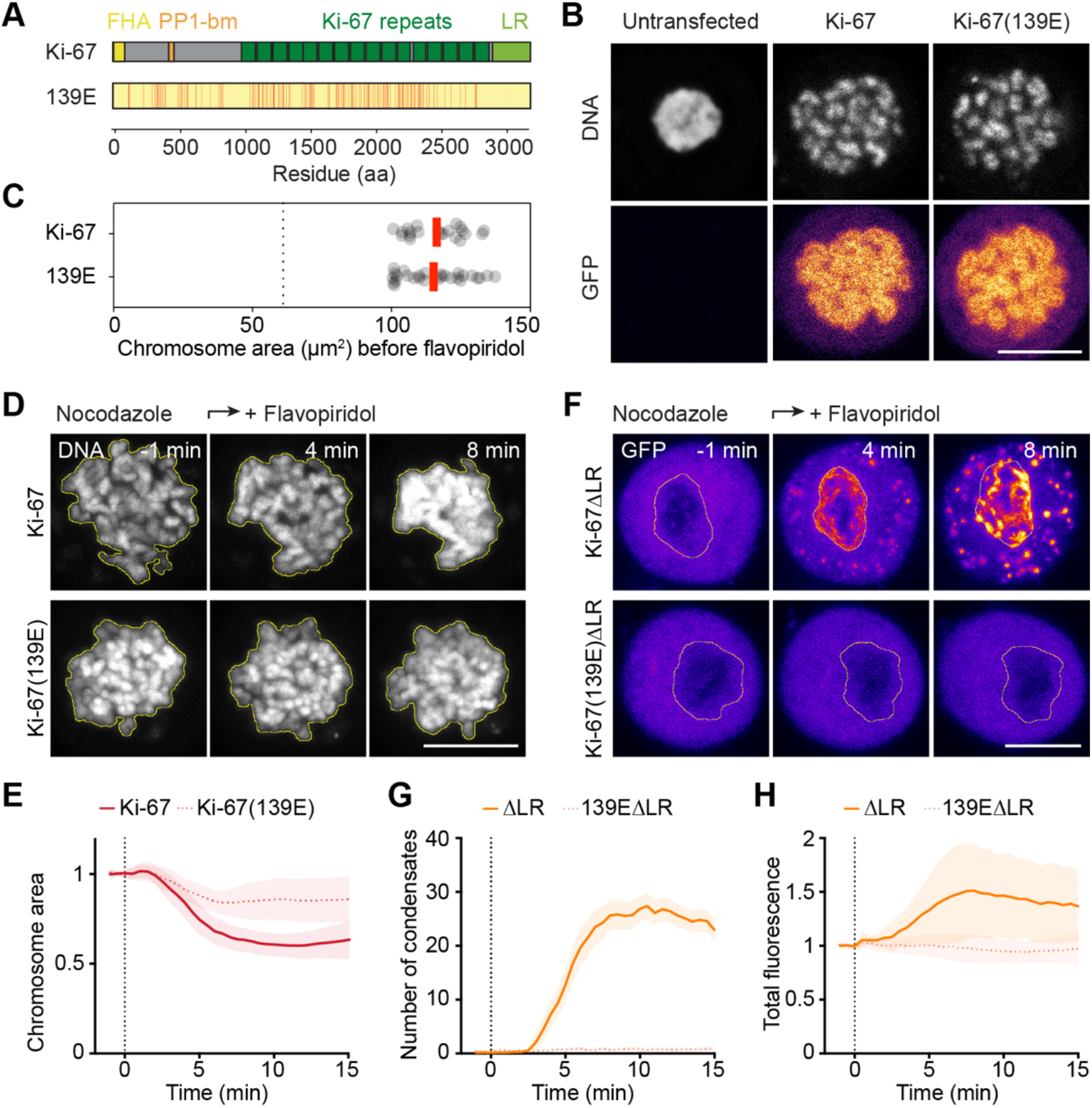
Ki-67 dephosphorylation controls Ki-67 phase separation, enrichment on chromosomes and chromosome clustering. (A) Scheme of Ki-67’s domain structure and position of 139 serine or threonine to glutamate substitutions to generate phosphomimetic Ki-67(139E) indicated by red lines. FHA = fork-head associated domain. PP1-bm = PP1 binding motif. LR = leucine/arginine-rich DNA-binding domain. (B) Ki-67(139E) effectively disperses chromosomes in early mitosis. Confocal microscopy images of Ki-67 KO cells untransfected or transiently expressing wild-type EGFP-Ki-67 or EGFP-Ki-67(139E) arrested by nocodazole in early mitosis. (C) Initial chromosome ensemble area (prior to flavopiridol addition) of Ki-67 KO cells transfected with either EGFP-Ki-67 or EGFP-Ki-67(139E) plotted in (E). Dashed vertical line indicate mean chromosome ensemble area of untransfected Ki-67 KO cells from Figure 1E for reference. (D) Time-lapse confocal microscopy images of HeLa Ki-67 KO cells expressing EGFP-Ki-67 or EGFP-Ki-67(139E) undergoing spindle-less mitotic exit. Yellow lines represent chromosomal regions used for quantifications in (E). (E) Quantification of chromosome ensemble area as in (D). n = 21 (Ki-67); n = 28 (Ki-67(139E)). (F) Time-lapse confocal microscopy images of HeLa Ki-67 KO cells expressing Ki-67ΔLR or Ki-67(139E)ΔLR undergoing spindle-less mitotic exit. White lines represent chromosomal regions used for quantifications in (F). (G) Quantification of number of cytoplasmic condensates as in (F). (H) Quantification of EGFP total fluorescence intensity in segmented chromosome areas as in (F). n = 23 (Ki-67ΔLR); n = 37 (Ki-67(139EΔLR)). Dashed vertical lines indicate flavopiridol addition (E, G, H). Time points relative to flavopiridol addition. Red bars indicate mean (C). Lines and shaded areas represent mean ± SD (E, H) or mean ± SEM (G). Single z-slices (B) or max. intensity z-projections (D, F) are shown. DNA was labelled with SiR-Hoechst. Scale bars, 10 µm (B, D, F).

Full-length Ki-67(139E) localized to the surface of mitotic chromosomes and restored chromosome dispersal during early mitosis in Ki-67 KO cells, which confirms that the mutations did not affect the surfactant function of Ki-67 (Figures 2B and 2C). Expression of Ki-67(139E) in Ki-67 KO cells did, in contrast, strongly reduce chromosome clustering upon exit from mitosis (Figures 2D and 2E; Video S3). This effect was presumably due to the failure of the mutant protein to enrich and form a condensed phase on the chromosome surface at mitotic exit. This hypothesis is supported by the finding that a phosphomimetic version that lacks the canonical LR DNA-binding domain (Ki-67(139E)ΔLR) neither accumulated on anaphase chromosomes nor formed condensates in the cytoplasm during exit from mitosis, in contrast to unmodified Ki-67ΔLR (Figures 2F–2H; Video S4).

### The basic patch of Ki-67 mediates chromosome clustering

To gain further insights into the mechanism of Ki-67 enrichment on chromosomes and phase separation, we mapped the domains of Ki-67 that are required for either of these two properties. Replacement of the LR domain by histone 2B as an alternative means to localize the protein to DNA had no effect on chromosome clustering (Figures S2A–S2D), which suggests that the LR domain makes no specific contribution to the clustering function beyond merely anchoring Ki-67 to chromosomes.

We next split the remaining protein sequence of Ki-67 into an N-terminal segment that precedes the Ki-67 repeat motifs (N terminus) and a C-terminal segment that contains the sixteen repeat motifs (C terminusΔLR; Figure 3A). Only the N-terminal segment bound to chromosomes with kinetics similar to Ki-67ΔLR and had the ability to phase separate (Figures 3B–3D; Video S5). Chromosomal enrichment of the C-terminal Ki-67 segment started, in contrast, only ∼5 min after induction of mitotic exit, at a timepoint when chromosome clustering had already largely completed (Figures 1A and 1B). Furthermore, the C-terminal segment did not form condensates, even when highly overexpressed (Figure S3). These results suggest that the ability of Ki-67 to phase separate is encoded in its N-terminal segment.

**Figure 3.**
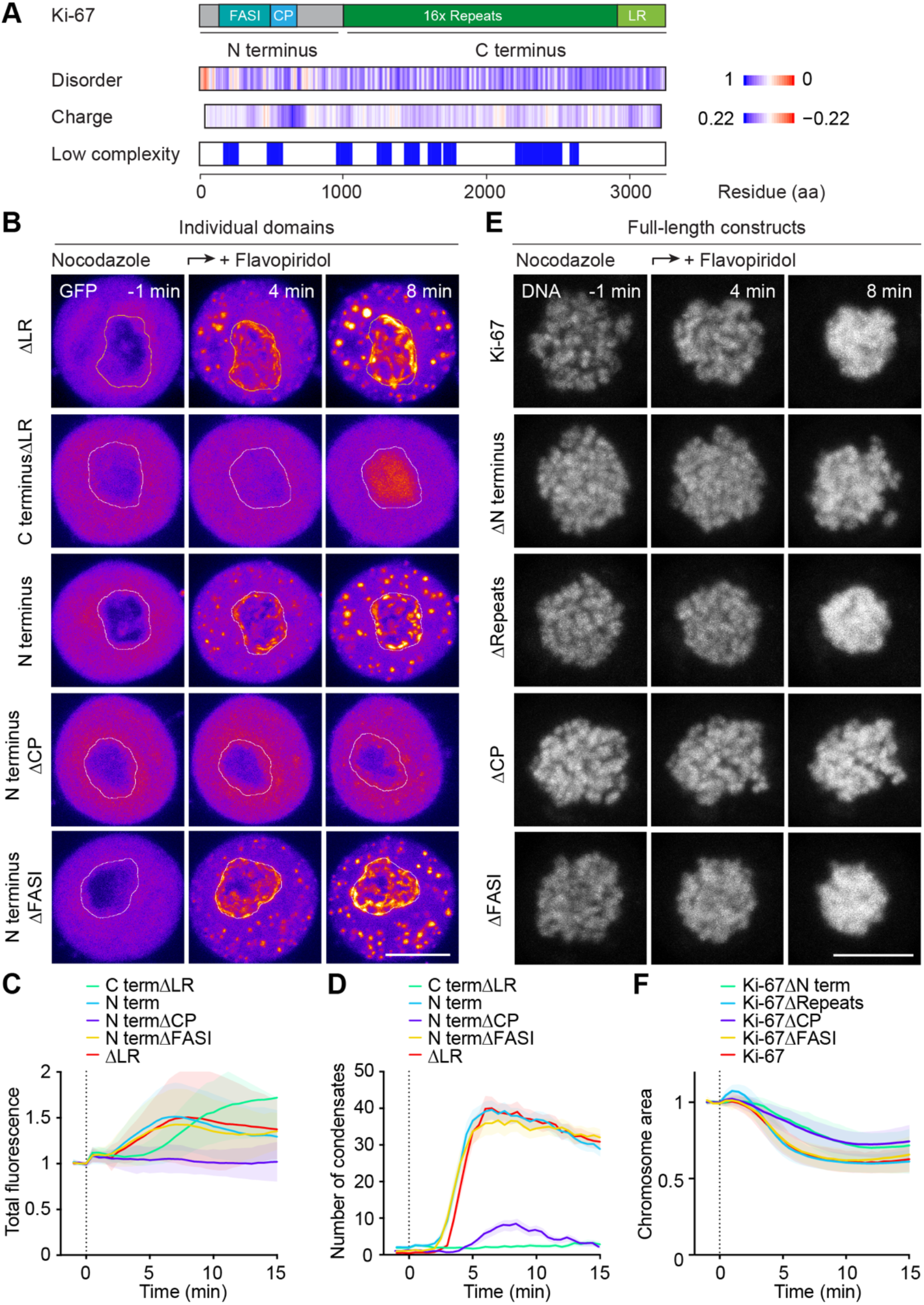
A positively charged patch (CP) within Ki-67 N terminus is responsible for Ki-67 phase separation, early-anaphase enrichment on chromosomes, and chromosome clustering. (A) Design of Ki-67 constructs. Disorder, charge and low complexity prediction of full-length Ki-67 based on IUPred3 (high disorder in blue, low disorder in red), EMBOSS charge webtool (sliding window 100, positive charge in blue, negative charge in red) and SEG webtool^21^ (sliding window size 45, low complexity regions in blue). CP = charged patch (186 aa). FASI = fragment absent in short isoform (360 aa). LR = leucin/arginine-rich DNA-binding domain. (B) EGFP-labelled domains of Ki-67 were transfected to HeLa Ki-67 KO cells and imaged during spindle-less mitotic exit. White lines represent chromosomal regions used for quantifications in (C). (C) Quantification of EGFP total fluorescence intensity in chromosomal regions as in (B). (D) Quantification of number of condensates in cytoplasm as in (B). n = 26 (ΔLR); n = 30 (C terminusΔLR); n = 33 (N terminus); n = 23 (N terminusΔCP); n = 29 (N terminusΔFASI). (E) Time-lapse microscopy images of Ki-67 KO cells expressing EGFP-labelled full-length Ki-67 or Ki-67 constructs with individual domain deletions as indicated undergoing spindle-less mitotic exit. DNA was labelled with SiR-Hoechst. (F) Quantification of chromosome ensemble area over time as in (E). n = 23 (Ki-67); n = 19 (Ki-67ΔN terminus); n = 26 (Ki-67ΔRepeats); n = 26 (Ki-67ΔCP); n = 22 (Ki-67ΔFASI). Dashed vertical lines indicate flavopiridol addition (C, D, F). Time points relative to flavopiridol addition. Lines and shaded areas represent mean ± SD (C, F) or mean ± SEM (D). Max. intensity z-projections are shown. Scale bars, 10 µm (B, E).

The N-terminal segment contains two low-complexity regions^21^: One that is present in only one of the two Ki-67 splice isoforms (Fragment Absent in the Short Isoform, FASI)^22^ and one that is located close to a positively charged patch (CP) of 186 amino acids (Figure 3A). Removal of the CP but not removal of the FASI region (Figures 3B–3D, and S3; Video S5) prevented enrichment on chromosomes and phase separation of the N-terminal segment during mitotic exit.

Consistent with the notion that the positively charged patch within the N terminus of Ki-67 is required for Ki-67 enrichment on chromosomes and phase separation, deletion of either the entire N-terminal segment or specifically of the CP from full-length Ki-67 considerably impaired chromosome clustering (Figures 3E, 3F, and S2F; Video S6). Deletion of the Ki-67 repeat domain or the FASI domain had, in contrast, no effect on the extent of chromosome clustering (Figures 3E, 3F, and S2F). Furthermore, a minimal construct comprised of CP-Repeats-LR clustered chromosomes to the same extent as wild-type Ki-67 (Figures S2E–S2G). Hence the N-terminal positively charged patch is necessary for chromosome clustering.

### High positive charge is the key feature of the charged patch

To further investigate how the CP drives chromosome clustering, we tested the relevance of the PP1-binding site and the high positive charge within the CP (Figures 4A and 4B). Both features are evolutionary conserved (^6^; Figure 4B). Mutation of the PP1-binding sequence RVSF to RASA^6^ resulted in a slight delay in Ki-67 enrichment on chromosomes, condensate formation, and chromosome clustering (Figure S4). These findings are consistent with the notion that dephosphorylation is critical for the switch in Ki-67 properties upon mitotic exit (Figure 2) and that PP1 contributes to its dephosphorylation.

**Figure 4.**
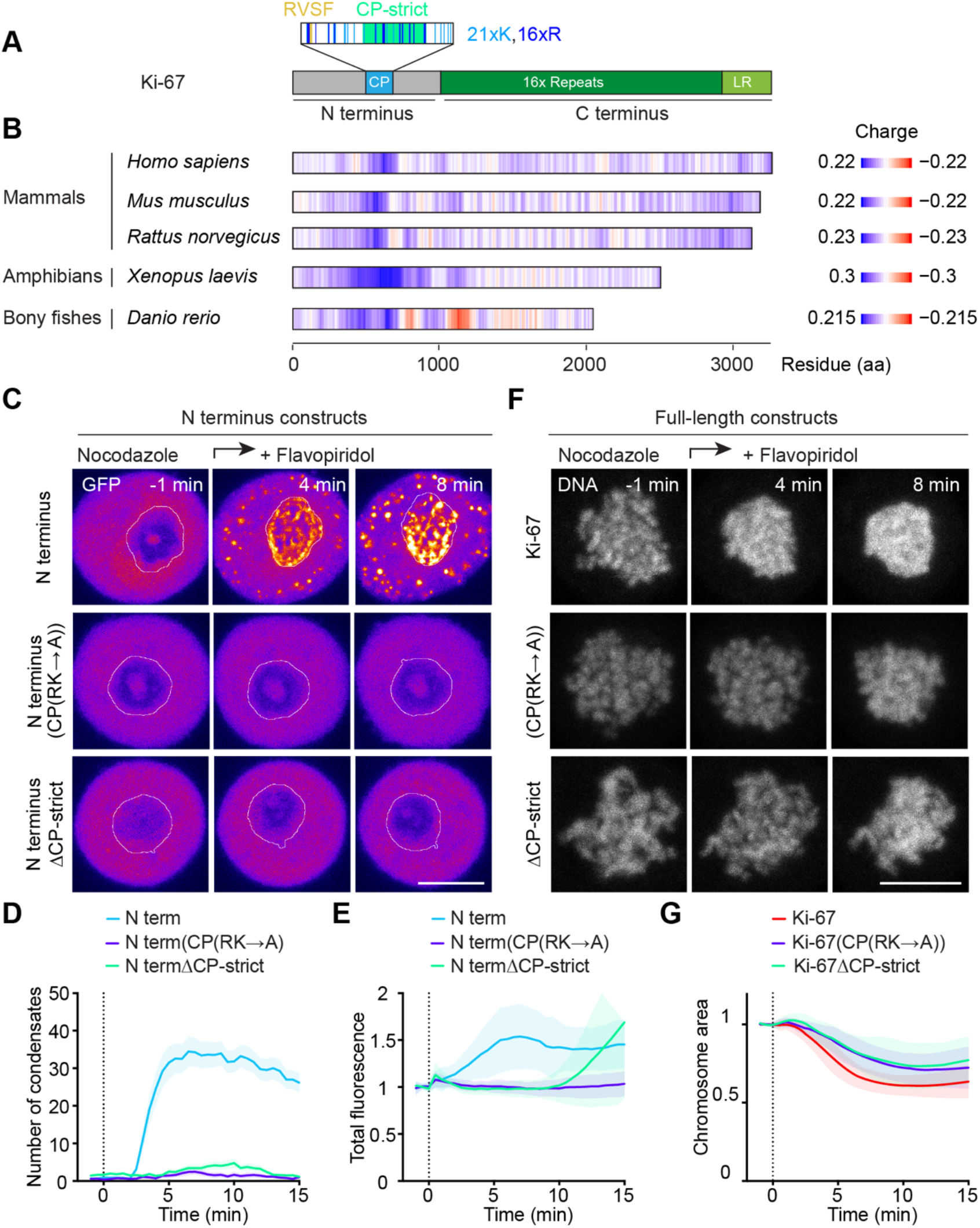
High positive charge density is highly conserved over predicted Ki-67 orthologues and is essential for chromosome clustering. (A) Schematic of Ki-67’s N-terminal segment. CP = Charged patch (186 aa) with lysine (K) and arginine (R) residues indicated. RVSF = Protein phosphatase binding motif mutated to RASA in Figure S4. CP-strict = Charge patch-strict (77 aa). (B) Heatmaps representing predicted Ki-67 orthologues using the EMBOSS charge webtool (sliding window of 100, positive charge in blue and negative charge in red). Notice the high positive charge density at the N terminus for all orthologues. NCBI reference sequences: *Homo sapiens* (NP_002408.3), *Mus musculus* (NP_001074586.2), *Rattus norvegicus* (NP_001258295.1), *Xenopus laevis* (NP_001128553.1), *Danio rerio* (NP_001264375.1). (C) Time-lapse microscopy images of spindle-less mitotic exit of HeLa Ki-67 KO cell transiently expressing different Ki-67 N terminus constructs tagged with EGFP. White lines represent chromosomal regions used for quantifications in (E). (D) Quantification of number of cytoplasmic condensates over time as in (C). (E) Quantification of EGFP total fluorescence intensity in segmented chromosome ensemble area over time as in (C). n = 22 (N terminus); n = 19 (N terminus(CP(RK®A))); n = 26 (N terminusΔCP-strict). (F) Time-lapse microscopy images of spindle-less mitotic exit of HeLa Ki-67 KO cell transiently expressing full-length Ki-67, Ki-67 constructs with individual domain deletions or mutations as indicated. DNA was labelled with SiR-Hoechst. (G) Quantification of chromosome ensemble area over time as in (F). n = 27 (Ki-67); n = 39 (Ki-67(CP(RK®A))); n = 25 (Ki-67ΔCP-strict). Dashed vertical lines indicate flavopiridol addition. Lines and shaded areas represent mean ± SD (E, G) or mean ± SEM (D). Max. intensity z-projections are shown. Scale bars, 10 µm (C, F).

Mutation of all 37 arginine and lysine residues within the CP to alanine (N terminus(CP(RK→A))) or deletion of only the section with the highest charge density (N terminusΔCP-strict) (Figure 4A) completely abolished condensate formation (Figures 4C, 4D, and S3; Video S5) and prevented, or considerably delayed, respectively, recruitment to chromosomes upon mitotic exit (Figure 4E). Consistent with this result, we found that full-length Ki-67 with charge-neutralizing mutations in the CP region (Ki-67(CP(RK→A))) or deletion of the CP section with the highest charge density (Ki-67ΔCP-strict) failed to cluster chromosomes to the same extent as wild-type Ki-67 (Figures 4F, 4G, and S2H; Video S6). We conclude that, during mitotic exit, enrichment of Ki-67 on chromosomes, Ki-67 phase separation, and chromosome clustering rely on the high density of positive charges within Ki-67 N terminus.

### Ki-67 phosphorylation counteracts chromosome clustering

To explore the role of phosphorylation in the function of the Ki-67 CP region, we restored the ten glutamate mutations of the phosphomimetic construct that are located in the CP region (Ki-67(129E); Figure 5A). In contrast to the original phosphomimetic Ki-67(139E) construct, the Ki-67(129E) version formed condensates upon mitotic exit (Figures 5A–5D) and supported chromosome clustering (Figures S2I, 5E, and 5F), albeit to a lesser extent than wild-type Ki-67. These findings suggest that some of the remaining phosphorylation sites, which are mostly located in the repeat sequences of Ki-67 (Figure 5A), also need to be dephosphorylated to complete chromosome clustering.

**Figure 5.**
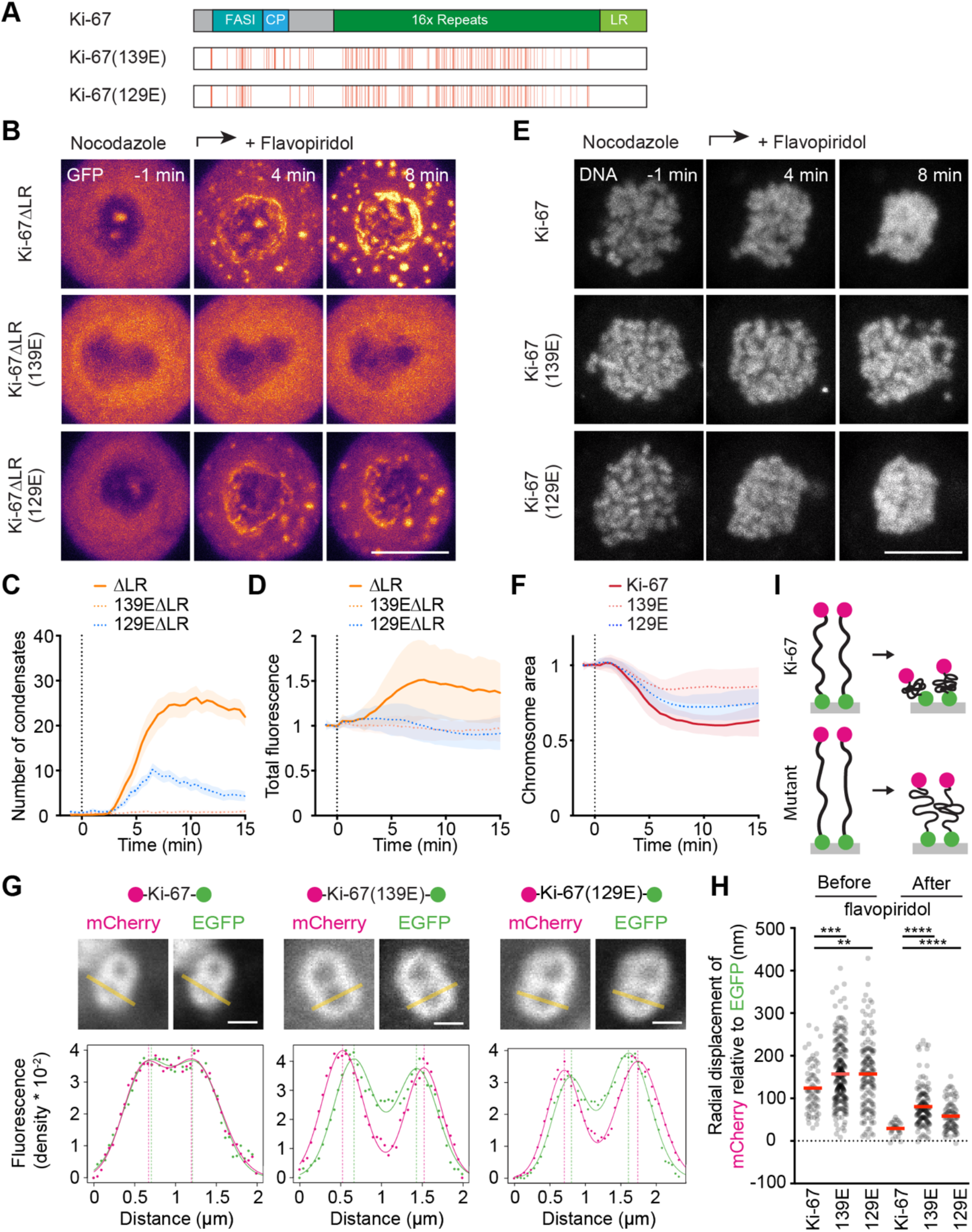
Complete Ki-67 dephosphorylation is required for full chromosome clustering. (A) Schematic of Ki-67 constructs tested. Position of 139 serine or threonine to glutamate (E) substitutions to generate phosphomimetic Ki-67(139E) marked by red lines. In Ki-67(129E) the 10 substitutions in the charged patch (CP) are reverted back to wild-type. (B) Time-lapse microscopy images of Ki-67 KO cells expressing EGFP-labelled Ki-67ΔLR constructs as indicated undergoing spindle-less mitotic exit. (C) Quantification of number of cytoplasmic condensates over time as in (B). (D) Quantification of EGFP total fluorescence intensity in chromosome ensemble area over time as in (B). Ki-67ΔLR and Ki-67(139E)ΔLR for reference as in Figure 2G and 2H; n = 31 (Ki-67(129E)ΔLR). (E) Time-lapse microscopy images of Ki-67 KO cells expressing full-length EGFP-labelled Ki-67 constructs as indicated undergoing spindle-less mitotic exit. DNA was labelled with SiR-Hoechst. (F) Quantification of chromosome ensemble area over time as in (E). Ki-67 and Ki-67(139E) for reference as in Figure 2E; n = 22 (Ki-67(129E)). (G) Wild-type Ki-67, Ki-67(139E) or Ki-67(129E) on the surface of mitotic chromosomes (taxol arrest) before and after flavopiridol addition. Constructs were labelled with mCherry at the N terminus and EGFP at the C terminus and expressed in HeLa Ki-67 KO cells (top). Images (middle) depict sister chromatid pairs oriented perpendicular to the imaging plane after flavopiridol addition. Yellow lines indicate measurements for the relative positions of mCherry and EGFP along the axis perpendicular to the chromosome surface (bottom). Sum of two Gaussian functions (lines) were fitted separately to the fluorescence densities (dots) along the line profiles of each fluorescent channel in order to determine peak positions (dashed lines). (H) Relative molecular extensions calculated from mCherry and EGFP radial displacements (mCherry peak-to-peak distance minus EGFP peak-to-peak distance /2) considering line profile measurements as in (G). Chromosome measurements before flavopiridol: n = 72 (Ki-67); n = 258 (Ki-67(139E)); n = 160 (Ki-67(129E)). Chromosome measurements after flavopiridol: n = 29 (Ki-67); n = 152 (Ki-67(139E)); n = 91 (Ki-67(129E)). Significance was tested by an unpaired t-test. (I) Model of molecular organisation of wild-type Ki-67 or Ki-67 mutants on the surface of chromosomes during early mitosis and during mitotic exit. Dashed vertical lines indicate flavopiridol addition (C, D, F). Lines and shaded areas represent mean ± SEM (C) or mean ± SD (D, F). Red bars indicate mean (H). Max. intensity z-projections (B, E) or single z-slices (G) are shown. Scale bars, 10 µm (B, E) or 1 µm (G).

We considered that dephosphorylation of these sites might be required to collapse the Ki-67 extended structure and allow complete chromosome clustering. To test this hypothesis, we measured the extension of the protein at the chromosome surface by utilizing Ki-67 constructs tagged with mCherry and EGFP at the N and C terminus, respectively. While the wild-type version decreased its distance between the fluorophores at mitotic exit to 29 nm (± 15 nm), the phosphomimetic Ki-67 mutants remained significantly longer in height (80 ± 46 nm for Ki-67(139E), 58 ± 34 nm for Ki-67(129E); mean ± SD), despite similar (139E) or slightly lower (129E) expression levels (Figures S2J and 5G–5I). This suggests that complete Ki-67 dephosphorylation is required for the full collapse of its molecular structure. Phosphorylation on the other hand promotes extension of the protein in early mitosis since the height of the phosphomimetic Ki-67 mutants were significantly larger. In summary, we conclude that Ki-67 phosphorylation counteracts chromosome clustering in two ways; by inhibiting its collapse and by neutralizing the positively charged patch.

### RNA is critical for chromosome clustering

How does the high density of positive charges mediate Ki-67 phase separation? Since RNA has been suggested as a universal factor that regulates phase separation^12,23,24^ and since RNA localizes to the chromosome surface^7,25^, we co-imaged RNA and Ki-67 in cells undergoing mitosis. The localization pattern of both molecule classes largely overlapped on the mitotic chromosome surface and in the cytoplasmic Ki-67 condensates that formed during anaphase (Figure 6A). RNA binding to chromosomes during metaphase and its increase during anaphase were strongly reduced in Ki-67 KO cells (Figure 6B), resulting in RNA accumulation in large cytoplasmic foci (Figure 6A).

**Figure 6.**
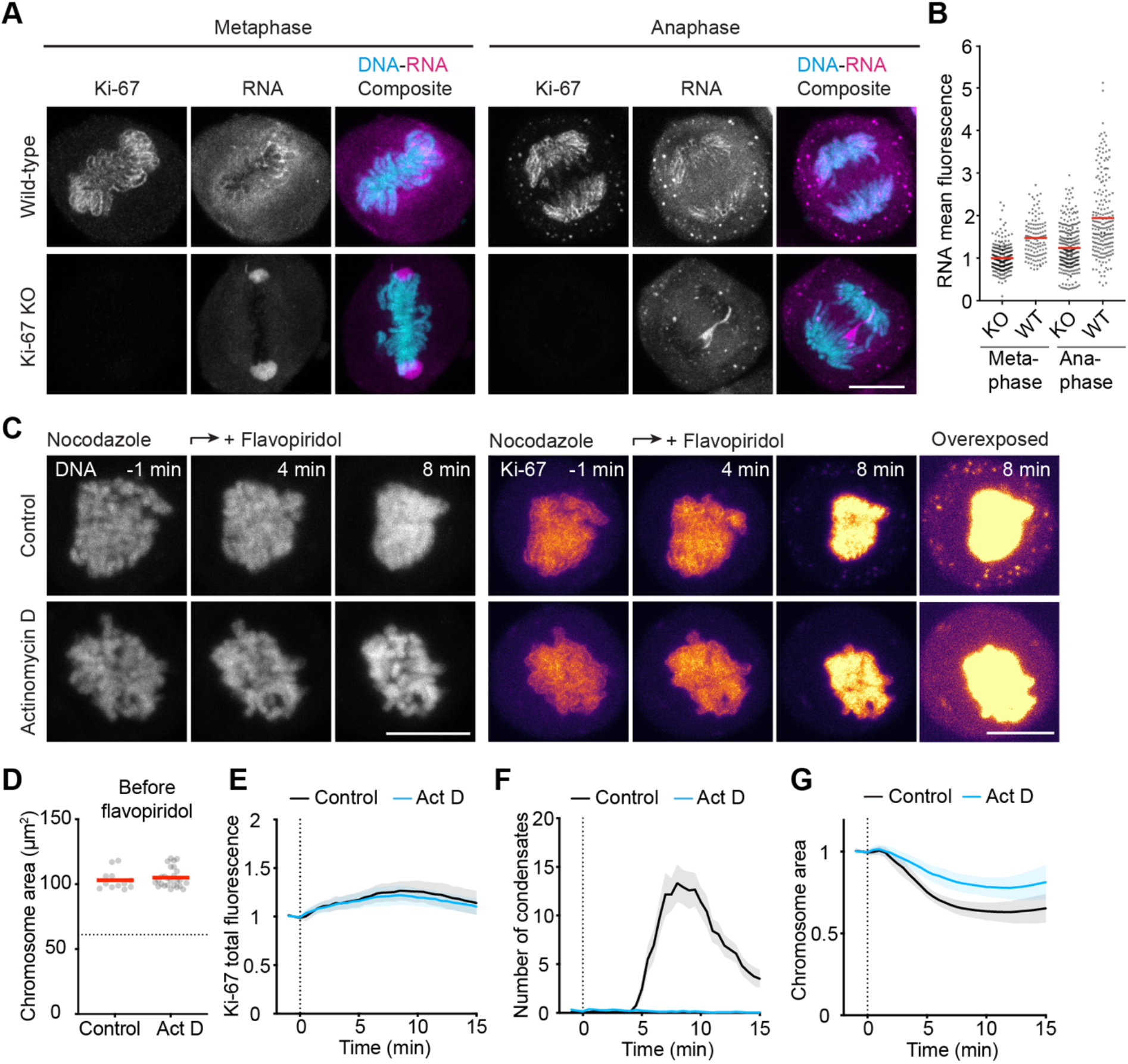
RNA is critical for chromosome clustering. (A) Immunofluorescence images of wild-type (WT) and Ki-67 KO cells (KO) undergoing metaphase (left) or anaphase (right). (B) Quantification of mean fluorescence intensity of EU-labelled RNA in DAPI-labelled chromosomes as in (A). Individual sets of chromosomes are plotted. n = 246 (metaphase KO); n = 118 (metaphase WT); n = 273 (anaphase KO); n = 198 (anaphase WT). (C) Time-lapse microscopy images of EGFP-Ki-67 knock-in cells either untreated (Control) or treated with actinomycin D [5 nM] (Act D) undergoing spindle-less mitotic exit. DNA was labelled with SiR-Hoechst. (D) Initial chromosome ensemble area (prior to flavopiridol addition) of EGFP-Ki-67 knock-in cells either untreated (Control) or ActD treated plotted in (G). Dashed horizontal line indicate mean chromosome ensemble area of Ki-67 KO cells from Figure 1E for reference. (E) Quantification of EGFP total fluorescence intensity in segmented chromosome area as in (C). (F) Quantification of number of cytoplasmic condensates over time as in (C). (G) Quantification of chromosome ensemble area over time as in (C). n = 12 (Control); n = 26 (Act D). Dashed vertical lines indicate flavopiridol addition (E–G). Red bars indicate mean (B, D). Lines and shaded areas represent mean ± SD (E, G) or mean ± SEM (F). Max. intensity z-projections are shown. Scale bars, 10 µm (A, C).

To test whether RNA plays a role in Ki-67 phase separation and chromosome clustering, we used endogenously tagged Ki-67 cells and depleted their ribosomal RNA (rRNA) which is the most abundant class of cellular RNAs and has been reported to bind Ki-67 in its precursor form^7,26^. Reduction of chromosomal rRNA levels by addition of the RNA polymerase I inhibitor actinomycin D (Act D) as verified by RNA-FISH (Figures S5A and S5B) neither affected chromosome dispersal upon mitotic entry nor Ki-67 enrichment during mitotic exit (Figures 6C–6E). However, it considerably reduced Ki-67 condensate formation (Figure 6F) and chromosome clustering (Figure 6G; Video S7). To ensure these effects are not due to side-effects of Act D, we confirmed these results with a different RNA polymerase I inhibitor, BMH-21, and obtained consistent results in an endogenously mCherry tagged Ki-67 cell line (Figures S5C–S5F) as well as in wild-type cells (Figures S5G–S5I). These data suggest that RNA-mediated phase separation of Ki-67, but not its enrichment on chromosomes during anaphase, is key to chromosome clustering.

### RNA is required for chromosome clustering in coarse-grained simulations

To further test this hypothesis, we performed molecular dynamics simulations using a minimal coarse-grained model. In these simulations, Ki-67 and RNA were represented as chains of charged beads connected by springs. To model Ki-67 dephosphorylation during mitotic exit, we assigned a charge that corresponds to 150 phosphorylation sites to each Ki-67 molecule and increased the charge as the simulation progressed until the molecule reached the net charge of fully dephosphorylated Ki-67 (Figure 7A).

**Figure 7.**
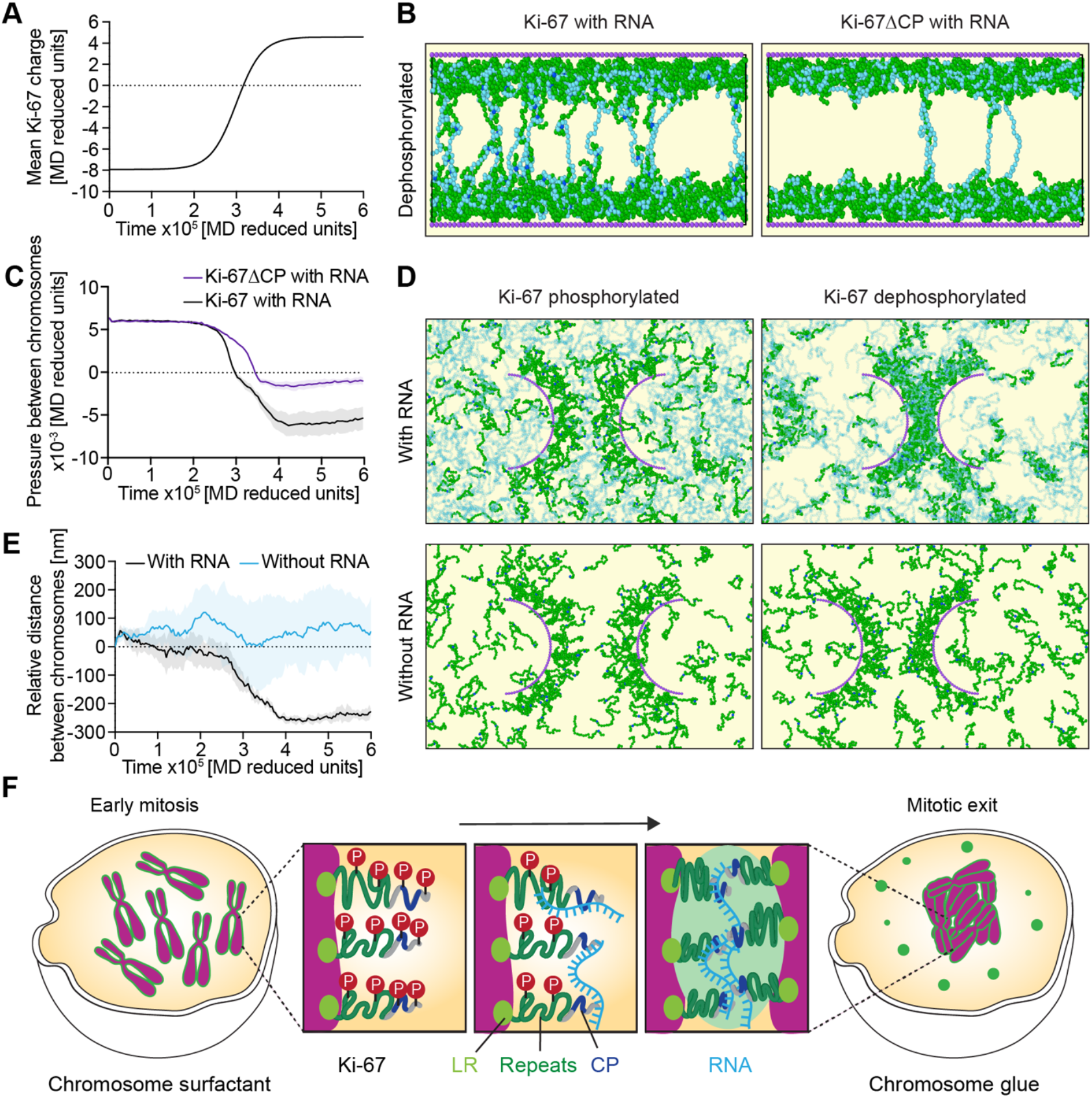
RNA is required for chromosome clustering in coarse-grained simulations. (A) Variation of mean Ki-67 charge per bead as a function of simulation time mimicking the switch from 150 phosphosites to fully dephosphorylated Ki-67. (B, C) Simulations of forces acting between two chromosome surfaces as the charge of Ki-67 is gradually increased as in (A). The two chromosome surfaces are modelled as immobile surfaces (violet), with a constant slightly negative charge, decorated with bead-spring polymers representing Ki-67 molecules (green – CP in dark blue) and constant highly negatively charged RNA molecules (light blue). (B) Simulation snapshots at a time point of fully dephosphorylated full-length Ki-67 with RNA (left) and Ki-67ΔCP with RNA (right). (C) Pressure (force per unit area) between chromosome surfaces as a function of simulation time using the gradual Ki-67 charge increase as in (A). n = 10 (Ki-67); n = 10 (Ki-67ΔCP). (D, E) Simulation of chromosome clustering as the charge of Ki-67 is gradually increased as in (A). The two chromosomes are modelled as half cylinders (violet) decorated with bead-spring polymers representing Ki-67 molecules (green – CP in dark blue). The chromosome surface has a constant, slightly negative charge, whereas the RNA molecules have a constant, highly negative charge. RNA molecules (light blue) are shown as semi-transparent for clarity. (D) Snapshots (top view) for Ki-67 charges corresponding to phosphorylated Ki-67 (early mitosis, left) or dephosphorylated Ki-67 (mitotic exit, right). (E) Relative distance (subtracting initial distance) between the centers of mass of the two chromosomes as a function of simulation time using the gradual Ki-67 charge increase as in (A). n = 5 (with RNA); n = 5 (without RNA). (F) Working model: During early mitosis, Ki-67 is heavily phosphorylated and extended, allowing dispersal of chromosomes. During mitotic exit, gradual dephosphorylation occurs, triggering both Ki-67’s charged patch (CP) to phase separate with RNA and the collapse of Ki-67, promoting coalescence of neighbouring chromosome surfaces and concomitant clustering of chromosomes. Dashed horizontal lines (A, C, E) are drawn at y = 0 for reference. Time points relative to simulation time (A, C, E). Lines and shaded areas represent mean ± SD (C, E).

Firstly, to simulate the forces between two chromosome surfaces, we modelled chromosomes as immobile surfaces decorated with grafted Ki-67 molecules and added freely diffusing RNA molecules (Figure 7B). When Ki-67 was in its phosphorylated state, the pressure acting on the two surfaces (force per unit area, for details see Methods) was positive, indicating a repulsive force between the surfaces. As Ki-67 became increasingly dephosphorylated, the pressure between chromosome surfaces decreased until it reached a negative equilibrium value, indicating an attractive force (Figures 7A–7C). In this state, the attractive force was exerted by negatively charged RNA molecules that interacted primarily with the positively charged patch of Ki-67 molecules, since simulations with Ki-67ΔCP molecules led to a higher equilibrium pressure (Figures 7B and 7C). Consistent with the notion that RNA and Ki-67 are necessary for the generation of attractive forces between chromosomes, we observed near-zero pressure between chromosomes in simulations performed in the absence of either RNA or Ki-67 (Figures S6A and S6B).

Secondly, to simulate chromosome clustering, we modelled chromosomes as mobile half-cylinders grafted with Ki-67 molecules in an environment of freely diffusing Ki-67 and RNA molecules (Figure 7D). Consistent with our experimental data (Figures 1A, 2, and 6A), the distance between the two chromosomes in the simulations decreased with increasing dephosphorylation of Ki-67, until an equilibrium distance was reached at which the two chromosomes were separated by a layer of Ki-67 mixed with RNA (Figures 7D and 7E; Video S8). In addition, in the simulations free Ki-67 molecules formed condensates with RNA independently of the presence of chromosomes (Figures 7D and S6C). Both, condensate formation and chromosome clustering, depended on the presence of RNA (Figures 7D, 7E, and S6C; Video S9), which is consistent with our experimental data (Figures 6C–6G, and S5C–S5I). Overall, our coarse-grained simulations (i) qualitatively reproduce the results of our cell-based experiments, (ii) are consistent with our proposed physical mechanism, and (iii) highlight the importance of RNA in chromosome clustering.

## DISCUSSION

Our cell-based experiments and computational simulations support a model (Figure 7F) in which, during early mitosis, phosphorylated Ki-67 protrude from the chromosome surface towards the cytoplasm to mediate the dispersion of individual chromosomes by a surfactant-like mechanism^8^. Upon mitotic exit, Ki-67 dephosphorylation causes the collapse of its extended structure and exposes a conserved positively charged patch within its amino terminus. These changes in conformation and biophysical properties reduce the amphiphilicity of Ki-67 at the chromosome-cytoplasm interface and, concomitant with RNA recruitment, trigger Ki-67 phase separation. Ki-67 condensation at the surface of chromosomes promotes chromosome clustering and, together with the concerted collapse of Ki-67, drives the exclusion of cytoplasmic material (Figure S7) before nuclear envelope reformation (^11^; Figures S1K and S1L).

A recent publication reported that single or multiple Ki-67 repeats can undergo phase separation *in vitro,* which is enhanced by phosphorylation^13^. This finding, in combination with the dynamic turnover of Ki-67 on chromosomes measured by FRAP and its dissociation from chromosomes observed upon sodium acetate treatment, led to the suggestion that Ki-67 might form a liquid-like phase on the surface of chromosomes during early mitosis. However, how such a liquid layer could contribute to chromosome dispersion during early mitosis had not been addressed.

We suggest that the opposite is the case: When two surfaces coated with a liquid interact, coalescence of the liquid layers should generate forces that bring the respective surfaces together – a feature that is in line with chromosome clustering, rather than with chromosome dispersion. Our data suggests that Ki-67 forms a liquid-like coat around chromosomes exclusively during exit from mitosis, triggered by its dephosphorylation (Figures 1 and 2) and RNA interaction (Figure 6). This process in turn, promotes chromosome clustering, which effectively excludes cytoplasmic material (Figure S7). During early mitosis, phosphorylation of Ki-67 instead inhibits phase separation (Figure 2) and causes the molecule to extend perpendicularly on the chromosome surface (Figures 5G–5I). Considering that phosphorylated Ki-67 likely has a negative net charge (Table S3) the extension away from the chromatin surface might be the result of repulsion of negative charges of both, phosphorylated Ki-67 and chromatin repelling each other. This leads to the formation of an electrostatic and steric barrier consistent with a surfactant-like mechanism.

The difference between our cellular findings and these earlier *in vitro* data^13^, especially regarding the specific region of Ki-67 responsible for phase separation and the impact of phosphorylation, may be attributed to the lack of RNA in the *in vitro* assays as well as to the use of high concentrations of both protein (40 µM in most *in vitro* assays as opposed to 50 nM endogenous in the cytoplasm (Figure S1H)) and crowding agents.

On the other hand, our findings support the notion that Ki-67’s interaction with RNA is key to phase separation and chromosome clustering. Several studies suggested that Ki-67 interacts either directly or indirectly with RNA: Recent reports showed that Ki-67 depletion caused loss of newly synthesized RNAs from chromosomes in early mitosis^7^, that recombinantly expressed fragments of Ki-67 such as the fork-head associated (FHA) domain or three tandem repeats pulled down pre-rRNAs from total RNA^26^, and that the FHA domain interacted with the putative RNA-binding protein NIFK^27^. Since the latter interaction was dependent on mitotic phosphorylation, it is, however, unlikely that NIFK binding contributes to RNA recruitment during mitotic exit. Nevertheless, indirect recruitment through one or more of the more than 60 other proteins that are recruited to the chromosome surface by Ki-67 during mitosis^4,28,29^ remains a possibility. Yet, based on the results of our experiments and simulations, it is tempting to speculate that direct complementary electrostatic interactions between negatively charged RNA molecules and the Ki-67’s positively charged patch contribute to their phase separation, possibly via complex coacervation^30,31^.

Other recent studies have shown that natural or artificial DNA-bound condensates can generate forces that remodel chromatin *in vitro*^32–34^ or in cells^35^ to mediate compaction or exclusion of chromatin. Our data suggest that electrostatic interactions contribute to the generation of a condensed phase on the surface of mitotic chromosomes, generating large enough adhesive forces to facilitate their clustering. The force-generating interface of the Ki-67/RNA condensed phase could in physical terms be viewed as capillary forces, which are well-known in non-living soft matter systems^36^ and have been suggested to play a role in multicellular tissue organization^37,38^ and intracellular condensate remodeling^39^. While our data reveal how phase separation is coordinated with lipid membrane confinement to re-establish nuclear and cytoplasmic compartments, regulated protein surfactants^8,40,41^ and surface-adsorbed condensates^42^ might have implications at surfaces and interfaces of other membrane-less biomolecular condensates to generate mechanical forces and structure biomolecular condensates.

## Acknowledgements

We thank Daniel W. Gerlich for providing cell lines, the EMBL Advanced Light Microscopy Facility (ALMF) for support and Christian H. Haering and Thomas Quail for input on the manuscript. This work was supported by the German Research Foundation (DFG project number 402723784) and the Human Frontier Science Program (CDA00045/2019). A.H.-A. and A.B. have received PhD fellowships from the Boehringer Ingelheim Fonds; V.S. and A.Š were supported by the European Research Council (ERC) under the European Union’s Horizon 2020 research and innovation programme (Grant No. 802960); Y. H. was supported by a fellowship from the EMBL interdisciplinary Postdoc (EIPOD) program (Marie Sklodowska-Curie Actions, COFUND grant agreement 664726).

## Author contributions

A.H.-A. and S.C.-H. conceived the project. A.H.-A. designed, performed and analyzed all experiments. V.S. and A.Š. designed simulations. V.S. performed and analyzed simulations.

Y.H. performed initial experiments on RNA localization in mitosis and contributed to RNA FISH experiments. A.B. helped A.H.-A. with acquisition of FCS-calibrated imaging. S.C.-H. acquired funding and supervised the project. S.C.-H., A.H.-A., A.Š. and V.S. wrote the manuscript with feedback from J.E.

## Data and code availability

Raw microscopy data and Fiji codes will be deposited on the BioImage Archive and GitHub, respectively. The simulation code is already available on Github (https://github.com/valesori/chromosome_clustering). Plasmids will be deposited at Addgene and cell lines will be available from the corresponding author upon request.

## Methods

### Cell lines and cell culture

All cell lines were regularly verified negatively for mycoplasma contamination and listed in Table S1. All HeLa cell lines were derived from a HeLa *Kyoto* cell line described in ^43^. Cells were cultured in Dulbecco’s modified medium (DMEM; Gibco, 41965039) containing 10% (v/v) fetal bovine serum (FBS; Gibco, 10270106), 1% (v/v) penicillin-streptomycin (Sigma Aldrich, 15140122) and 1 mM sodium pyruvate (Gibco, 11360039). DNA was visualized by labelling with 100 nM SiR-Hoechst^44^ unless otherwise indicated. Live cell imaging was performed in FluoroBrite™ DMEM (Gibco, A1896701) containing 10% (v/v) FBS, 1% (v/v) penicillin-streptomycin, 1% (v/v) GlutaMAX (Gibco, 35050038) and 1 mM sodium pyruvate. Cells were grown in LabTek chambered coverglass (Thermo Scientific).

### Plasmid transfections

Transfections were done transiently using PEI transfection reagent (1 mg ml^-1^ stock, Polysciences, 24765-1), 4 µg of transfection reagent per 1 µg of plasmid, and incubated for 48 h before imaging.

### Generation of Ki-67 constructs

All constructs are listed in Table S2. Regions within Ki-67 (3256 total amino acids) were defined as follows: Ki-67’s N terminus: residue 1 to 1002; Ki-67’s C terminusΔLR (also called Repeats): residues 1003 to 2929; Ki-67’s LR domain: residues 2930 to 3256; Ki-67’s FASI: residue 136 to 495; Ki-67’s Charged Patch (CP): residue 496 to 681; Ki-67’s Charged Patch-strict (CP-strict): residue 571 to 647. The phosphomimetic version of Ki-67 (Ki-67(139E)) was generated by Gibson assembly of three gBlocks (IDT) into an IRESpuro2 backbone with N-terminal EGFP. Plasmids will be been deposited at Addgene.

### Inhibitors and stains

Cells were arrested in prometaphase by incubating them for 2 – 4 h in 200 ng ml^-1^ nocodazole (Sigma-Aldrich, 31430-18-9). Mitotic exit was induced by adding flavopiridol (Tocris Bioscience, 131740-09-5) to a final concentration of 20 µM. All drug treatments were performed for 3 – 5 hours prior to imaging with a final concentration of 5 nM of actinomycin D (Gibco, 11805-017) or 1 µM of BMH-21 (Sigma-Aldrich, 896705-16-1).

### RNA visualization by 5’EU labelling and Ki-67 immunofluorescence

HeLa wild-type cells and HeLa Ki-67 KO cells were co-seeded in single wells and synchronized by a double thymidine block. Specifically, cells were incubated for 24 h in medium containing 2 mM thymidine (Sigma-Aldrich, 50-89-5), followed by release in fresh medium. After 8 h, the medium was exchanged for 2 mM thymidine-containing medium and incubated for 16 h. Afterwards, the medium was exchanged for fresh medium and incubated for 6 h. Then, cells were incubated in medium containing 200 µM of 5-Ethynyl Uridine (Thermo Fisher Scientific, E10345) and after 3 h, cells were washed with PBS and fixed with 4% PFA (Merck, 50-00-0) for 15 min at room temperature. Afterwards, the fixed cells were washed twice with PBS and permeabilized with 0.2 % Triton X-100 (Roth, 9002-93-1) for 5 min at room temperature, followed by two washes with 3 % BSA (Sigma-Aldrich, 9048-46-8). For labeling, fixed cells were treated with Click-iT™ Plus (Invitrogen, C10643) reaction cocktail for 30 min in the dark, washed twice with 3 % BSA and incubated for 1 h with monoclonal rabbit anti-Ki-67 antibody (Abcam, ab16667, 1:1000) and washed three times with PBS. Alexa 594 goat anti-rabbit IgG (Invitrogen, A11037, 1:500) was used for visualization, incubated for 45 min and washed three times with PBS. Finally, DNA was incubated with DAPI (Thermo Fisher Scientific, 62248, 1:1000) for 5 min and washed once with PBS.

### Pre-ribosomal RNA visualization by RNA-FISH of ETS

Homozygous endogenously EGFP-labelled Ki-67 HeLa cells or HeLa Ki-67 KO cells were seeded and synchronized by a double thymidine block (see above). After 6 h of the second thymidine release, cells were either untreated or treated with actinomycin D or BMH-21. After 3 h, cells were fixed and permeabilized as previously described. Afterwards, cells were washed three times with PBS for 5 min, incubated with wash buffer (10 % formamide (Sigma-Aldrich, 75-12-7) in 2× saline sodium citrate (SSC; Merck, 6132-04-3)) for 5 min at room temperature. Then, the solution was exchanged to hybridization buffer (125 nM ETS1-1399 (Cy35’CGCTAGAGAAGGCTTTTCTC3’Cy3^45^), 10% formamide, 2× SSC, 10% dextran sulfate (Merck, S4030), 10 mM ribonucleoside-vanadyl complex (NEB, S1402S)) and incubated overnight at 37°C in a humidified chamber. Cells were gently washed twice with wash buffer in the dark for 15 min at 37°C. DNA was labelled with DAPI (Thermo Fisher Scientific, 62248, 1:1000) at room temperature for 5 min and washed twice with PBS.

### Microscopy

Imaging was performed on a customized confocal Zeiss LSM780 microscope, using × 40 or × 63, 1.4 NA, Oil DIC Plan-Apochromat objective (Zeiss), operated by ZEN 2011 software, and an incubator chamber (European Molecular Biology Laboratory (EMBL), Heidelberg, Germany) provided constant humidity and 37°C temperature with 5% CO_2_.

### Quantification of chromosome ensemble areas and protein enrichment

Maximum intensity projections (4 z-sections with 2 µm spacing recorded) of cells expressing EGFP- or mCherry-tagged proteins were analyzed using CellCognition Explorer^46^, where a primary segmentation mask in the chromatin channel was defined by applying a local adaptive threshold, and both channels quantified. Using R, the signals were normalized to the mean value before flavopiridol addition for each cell.

### Quantification of GEMs in chromosome ensemble area

Quantification of GEMs was done by segmenting the chromosome ensemble area in a single central z-slice (4 z-sections with 2 µm spacing were recorded). If bleaching was observed, a frame-wise exponential fitting using the Image J plug-in “Bleach correction” (htt://fiji.sc/Bleach_Correction) was performed before segmentation. The bleached-corrected chromatin channel of the central single z-slice was denoised using a Gaussian blur filter (σ = 2) and thresholded using ImageJ’s default method (a variation of the IsoData algorithm) and converted to a binary image. Next, the Fiji convex hull algorithm (https://blog.bham.ac.uk/intellimic/g-landini-software/) was applied to the binary image and used as a mask for the quantification of GEM fluorescence intensity in the chromosome ensemble area of the central single z-slices. All cells were normalized to the average GEM total fluorescence (in the chromosome ensemble area) before flavopiridol addition.

### Quantification of cytoplasmic condensate number and total condensates area

To only measure cytoplasmic Ki-67, a chromosome mask was generated based on SiR-Hoechst images and used to cancel any Ki-67 signal in that area. In detail, the chromosome area was segmented in maximum intensity z-projections of SiR-Hoechst images (4 z-sections with 2 µm spacing) which were denoised using Gaussian blur (σ = 2), thresholded using ImageJ’s default automated method and converted to a binary image. This mask was dilated with 20 iterations, to avoid any fluorescent signal originating from the chromosome ensemble area, and inverted resulting in 0 values for the chromosome mask. To quantify the number of condensates in the cytoplasm only, the inverted mask was multiplied with the GFP channel cancelling any GFP signal in the chromosome area. The GFP image was denoised using Gaussian blur (σ = 2) and the background was subtracted using the “subtract background” function with a 25-pixel rolling ball radius. The threshold was set to (0, 2500) for all N terminus and full-length mutants tested, and to (0, 2000) and (0, 1000) for quantification of endogenously tagged EGFP- and mCherry-Ki-67 cell lines, respectively. In addition, a frame-wise exponential fitting using the Image J plug-in “Bleach correction” (htt://fiji.sc/Bleach_Correction) was performed before condensate segmentation for mCherry-Ki-67 expressing cells. The respective image was transformed into a mask and the function “Analyze particles” was used to quantify the number of condensates and the sum of their areas per frame.

### Quantification of absolute Ki-67 concentration during mitotic exit

To quantify absolute concentrations of Ki-67 during mitotic exit in living cells, fluorescence correlation spectroscopy (FCS)-calibrated imaging was performed as in^47^. In brief, HeLa Kyoto cells were either untransfected (for estimation of background photon counts) or transfected with mEGFP (for FCS measurements of freely diffusing mEGFP) 24 h prior to imaging. HeLa EGFP-Ki-67 knock-in cells^8^ were seeded 48 h prior to imaging.

Acquisition was carried out on a Zeiss LSM880 using a C-Apochromat × 40, 1.20 NA, W Korr FCS M27 water-immersion objective. Confocal volume estimation was carried out by ten FCS-measurements of 1 min of 10 nM Atto488 (AD 488-21, ATTO-TEC) in double-distilled water. Of note, Atto488 carboxylic acid was used for the effective confocal volume measurement instead of the previously reported AF488-NHS^47^. Use of the Atto488 dye in combination with its reported diffusion coefficient of 400 µm^2^/s at 25°C (PicoQuant) typically results in larger effective confocal volume estimates and therefore lower absolute concentrations compared to AF488-NHS^47^. Background fluorescence and background photon counts were determined by FCS measurements targeted to both the nucleus and cytoplasm of untransfected HeLa Kyoto cells. Subsequently, repeated imaging and nuclear and cytoplasmic FCS measurements of HeLa Kyoto cells expressing a broad range of mEGFP were recorded. In combination with the estimated effective confocal volume, an experiment specific internal calibration factor was computed with which measured mEGFP fluorescence intensities could be converted into protein concentrations in any image that was acquired with the same imaging settings.

Mitotic exit movies of unsynchronized EGFP-Ki-67 knock-in cells were acquired by automated detection of metaphase cells in large fields of view using low-resolution imaging of the SiR-Hoechst channel and ilastik-based image segmentation and classification^48^, followed by high resolution imaging every minute for 40 min.

Quantification of fluorescence intensities was performed in single-Z slices using Fiji. Cytoplasmic condensates were segmented as described above. For chromosomes, a mask was generated based on SiR-Hoechst images and used to measure the Ki-67 signal in that area. Specifically, the chromosome area was segmented, denoised using Gaussian blur (σ = 2), thresholded using ImageJ’s default automated method, converted to a binary image, and dilated with 5 iterations. Next, the GFP image was denoised using Gaussian blur (σ = 2) and background subtracted with 25-pixel rolling ball radius. Then, thresholded to (0, 5000) and the respective images were transformed into a mask and the function “Analyze particles” was used to quantify the intensities. For quantifying the cytoplasmic fluorescence intensity, a circular ROI (1.5 µm of diameter), was used to follow the fluorescence of the cytoplasm over time.

### Fluorescence recovery after photobleaching (FRAP) experiments

EGFP-labelled Ki-67ΔLR was transiently transfected to Ki-67 KO cells and spindle-less mitotic exit was induced as previously described. Six min after flavopiridol addition, a pre-bleach time series was acquired for at least 9 s. A circular ROI (diameter between 0.5 to 1.0 µm) was used for partially bleaching condensates of at least 1 µm in diameter. To capture recovery, time series of at least 60 s were acquired with 0.5 s intervals. Similarly, for addressing mobility of Ki-67’s chromosome-bound pool, a circular ROI (diameter of 2 µm) was used for bleaching EGFP-Ki-67 knock-in cells chromosomes, previous to flavopiridol addition and after 3 min. Both conditions, considered pre-bleach time series for at least 9 s and the recovery of fluorescence was followed for at least 3 min with 0.5 s intervals.

### Quantification of Ki-67’s molecular extension in early mitosis and mitotic exit

To measure Ki-67 molecular extension at the chromosome surface, both termini of Ki-67 were tagged with different fluorescent proteins and Gaussian functions were fitted to their line profiles as described in^4,5^. Ki-67 KO cells were transfected with either mCherry-Ki-67-EGFP, mCherry-Ki-67(139E)-EGFP or mCherry-Ki-67(129E)-EGFP, arrested in medium containing

0.5 µM taxol (Sigma, T1912) for 1 h and recorded before and 5 – 8 min after flavopiridol addition. Only chromosomes with their arms oriented perpendicular to the imaging plane were analyzed, by measuring line profiles of red and green channels that sectioned a single sister chromatid. After background subtraction and normalization to total fluorescence intensity, the sum of two Gaussian functions was fitted and the distance between the Gaussian center positions was measured separately for green and red fluorescent channels using R. This value represents the sum of both chromatic shifts perpendicular to the chromosome surface, and the molecular extension of Ki-67 can be calculated by dividing this value by two.

### Coarse-grained model of chromosome surfaces, Ki-67 and RNA

To model the dispersion and clustering of chromosomes in the presence of Ki-67 and RNA, we perform *NV T* (constant number of particles, volume and temperature) molecular dynamics (MD) simulations of a coarse-grained model. Chromosome surfaces are modeled as rigid bodies, generated by arranging spherical beads of diameter *σ* on a surface (either a surface or a half cylinder, see below). In experimental units, *σ* ≃ 16 nm (see discussion about units conversion below). The bead surface number density is *ρ_s_* = *σ^−^*^2^.

Ki-67 molecules can be either grafted to the chromosome surfaces or free (the latter molecules modelling cytoplasmic Ki-67). Each molecule is modeled as a bead-spring polymer chain of *N*_Ki67_ = 24 beads (bead diameter = *σ*). The grafted molecules are bound to the chromosome surface by a finite extensible nonlinear elastic (FENE) bonds (see Eq. (4) below), *i.e.*, the LR domain and its interaction with the chromosome is not modeled explicitly. RNA molecules are also modeled as bead-spring polymer chains of *N*_RNA_= 100 beads (bead diameter = *σ*) which can freely move in the simulation box. To investigate the role of RNA in chromosome clustering, we have also considered systems without RNA.

The interaction between all the beads (belonging to the chromosomes, to the Ki-67 and to the RNA) is modeled by an excluded volume component *E*_exc.vol_ (truncated and shifted Lennard-Jones potential), which models steric interactions, and a screened electrostatic component *E*_el_. The total interaction potential is thus

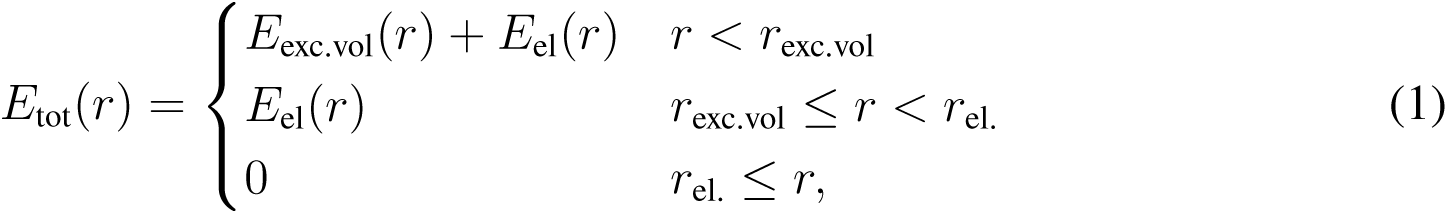

where *r* is the distance between the centers of the beads and the two interaction potentials are given by

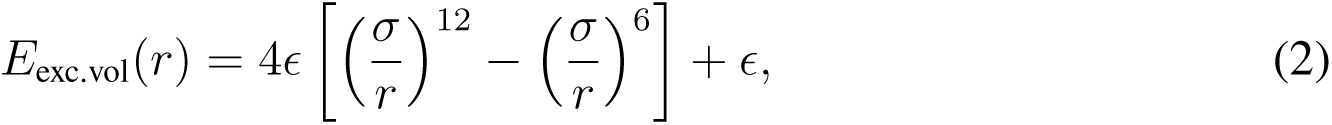

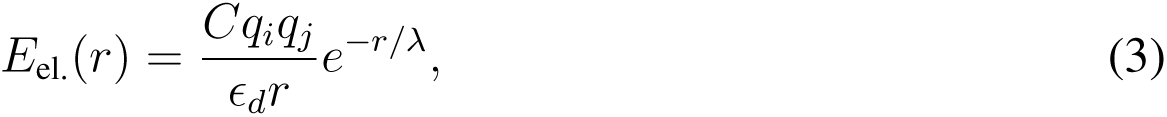

and *r*_exc.vol_ = 2^1^*^/^*^6^*σ*, *r*_el._ = 3*σ* are the cutoff lengths of the two potentials. In Eq. (2), *ɛ* is the interaction energy. In Eq. (3), *C* is a unit conversion constant, *q_i_, q_j_* are the charges of the two interacting particles, *ɛ_d_* = 5.0 is the dielectric constant and *λ* = *σ/*2 is the Debye screening length. In what follows, we use MD reduced units (*ɛ* = 1, *σ* = 1, *k_B_* = 1 and *m* = 1, with *m* the mass of the beads comprising the Ki-67 and RNA polymers). All the other quantities will be expressed in these MD reduced units unless explicitly stated.

In addition to the two terms of Eqs. (2) and (3), bounded neighbors in the same polymer chain interact via a finite extensible nonlinear elastic (FENE) potential:

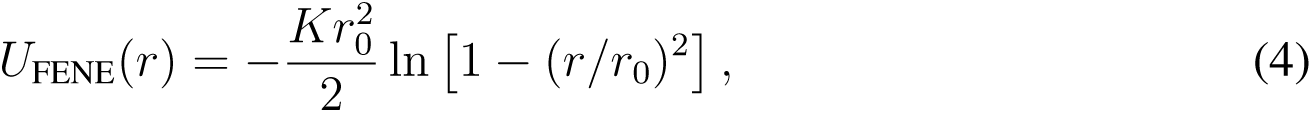

with *K* = 30 and *r*_0_ = 1.5 (Kremer-Grest model ^50^). These values are chosen in such a way to prevent chain crossing. In addition to the FENE potentials, bonded triplets interact through a bending potential, to simulate semiflexible polymers ^51^:

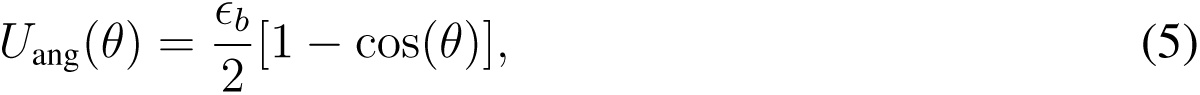

where *θ* is the triplet angle and *ɛ_b_* = 2 is the bending stiffness. The solvent is simulated implicitly using a Langevin thermostat ^52^. The thermostat also insures that the temperature *T* is kept constant during the simulation at *T* = 1, which corresponds to 37*^◦^* C in this case. The viscous friction that each bead experiences is *ζ* = 10. The simulations are carried out using the LAMMPS software ^49^, and time integration is performed using the velocity Verlet algorithm, with time step *δt* = 2 × 10*^−^*^3^.

### Charge distribution of Ki-67 and RNA molecules

To each bead comprising the RNA molecules we assign the same negative charge *q*_RNA_ = − 15, so that RNA polymers are uniformly charged. The total charge of an RNA molecule *N*_RNA_*q*_RNA_ = –1500. We have also performed simulations with *q*_RNA_ = –10, finding the same qualitative results (not shown). The beads comprising the surfaces of the two chromosomes have a weak negative charge *q*_chr_ = –0.5. The Ki-67 polymers are also uniformly charged (each bead having charge *q*_Ki67_), with the exception of the fifth bead from the polymer’s end, which is given a higher charge *q*_CP_. This mimics the charged patch (CP) experimentally observed in Ki-67 molecules. The charges *q*_Ki67_ and *q*_CP_ are chosen to mimic the charge state of Ki-67 during early mitosis at the start of the simulation (*i.e.*, at time *t* = 0). In particular, we consider the theoretical charge distribution of a Ki-67 molecule with 150 phosphorylated amino-acids (thus adding a total charge –300 in atomic units – see below for more details on how the theoretical distribution is calculated). Thus, the initial charge of the Ki-67 beads (excluding the CP) is *q*_Ki67_(0) = –8.54 and the initial charge of the CP is *q*_CP_(0) = 6.17 [total charge *Q*_Ki67_(0) = 23*q*_Ki67_ + *q*_CP_ = − 190.2].

To simulate the gradual switching of Ki-67 from the phosphorylated to the dephosphorylated state, the charge of both the CP and of the rest of the chain is increased according to the following sigmoidal function:

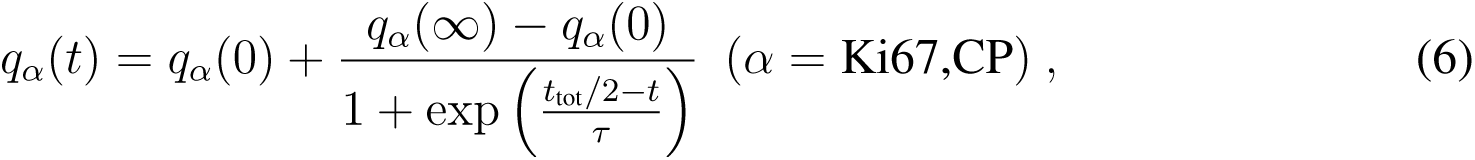

where *t*_tot_ = 6 × 10^5^ is the total simulation in MD reduced units, and *τ* = 3 × 10^4^ is a typical time for the change variation. With this choice, at the end of the simulation the charge distribution of the Ki-67 molecules corresponds to the mitotic exit (fully dephosphorylated) state, *i.e.*, *q*_Ki67_(*t*_tot_) ≃ *q*_Ki67_(∞) = 3.73 and *q*_CP_(*t*_tot_) ≃ *q*_CP_(∞) = 24.15 [total charge *Q*_Ki67_(∞) = 109.8].

### Measurement of the force per unit area acting between chromosome surfaces

To precisely quantify the force acting between the chromosomes, we perform simulations in which the Ki-67 molecules are grafted to surfaces, representing the surfaces of the chromosomes. As the focus of these simulations is not to study the formation of condensates of Ki-67 and RNA in solution, we do not include free (cytoplasmic) Ki-67 molecules. The two square surfaces (area *A* = 4.9 10^2^) are kept fixed in space at a constant distance *L_z_* = 30 from each other, with the Ki-67 brushes facing each other. We graft to each surface 100 Ki-67 molecules (total number of Ki-67 molecules in the system *n*_Ki67_ = 200), which are regularly arranged on a square lattice and bound to the surface by FENE bonds (Eq. (4)). The surface density of the grafted Ki-67 is *ρ_g_* = 2.0 10*^−^*^2^, or approximately 78 *µ*m*^−^*^2^ in experimental units. We additionally simulate *n*_RNA_ = 24 free RNA polymers of length 100 in the space comprised between the two grafted surfaces. Periodic boundary conditions are applied in the *x* and *y* directions, which are parallel to the surfaces.

The system is equilibrated for a time 10^5^ using only the excluded volume interactions before the production run with the full interactions. During production (duration *t*_tot_ = 6 × 10^5^), the charge of the Ki-67 molecules is changed according to Eq. (6) to model dephosphorylation. During the production run, we measure the *zz*-component of the stress tensor, *σ_zz_*, evaluated using the *compute pressure* LAMMPS command. This quantity has the dimensions of a pressure and corresponds to the force per unit area acting on the two surfaces. If its value is negative, there is an attractive force between the two surfaces; if it is positive, the force is repulsive. To assess the role of the CP in determining the strength of the interaction between the two surfaces, we additionally perform simulations in which the CP is completely removed (the Ki-67 polymer thus comprising of 23 instead of 24 beads). For each set of parameters, we perform ten independent simulation runs (equilibration and production) and average the results over these simulations. In addition to these simulations, we also perform some in which RNA or Ki-67 is removed, in order to measure the force between the two surfaces in the absence of one of these two molecules. For these simulations, we performed five independent runs.

### Simulation of chromosome clustering

To model chromosome clustering triggered by the dephosphorylation of Ki-67 and mediated by RNA, we model the chromosome sections as rigid half cylinders with radius *R* = 17.2 and height *h* = 30. The two half cylinders face each other at the start of the simulation and are constrained to only undergo translational motion with respect to one another, along the *x* axis of the box. The mass of the beads comprising the chromosomes is set to *m* = 0.1, so that the total mass of a chromosomes is *M*_chr_ = 185.5. This is done in order to facilitate the movement of the chromosomes. A larger mass would lead qualitatively to the same dynamics, but would require a longer simulation time to observe separation/clustering of the chromosomes. The simulations are carried out in a rectangular cuboid simulation box with edges *L_x_* = 120 and *L_z_* = *L_y_* = 75, and periodic boundary conditions in all three spatial directions.

The Ki-67 brush is modeled by grafting *n*_Ki67_ = 80 bead-spring polymer chains of *N*_Ki67_ = 24 beads on the surface of the chromosomes. These chains are regularly arranged on a square lattice on the chromosomes’ surface and bound to it by a finite extensible nonlinear elastic (FENE) bonds. Cytoplasmic Ki-67 is modeled by 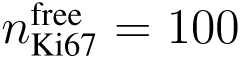 identical bead-spring polymers (without the LR domain) which are not bound to the chromosomes’ surface and are free to move in the simulation box. Due to the different geometry, for simplicity we choose a slightly higher surface density of the grafted Ki-67 than the one used for the simulations with the fixed surfaces, namely *ρ_g_* = 2.5 10*^−^*^2^, or approximately 98 *µ*m*^−^*^2^ in experimental units. Finally, we also model RNA as *n*_RNA_ = 100 bead-spring polymer chains of *N*_RNA_ = 100 beads (bead diameter = *σ*); the RNA molecules can also freely move in the simulation box. To investigate the role of RNA in chromosome clustering, we have also considered systems without RNA.

In order to study the effect of the Ki-67 charge distribution on the interaction between chromosomes, we initially place them facing each other with their surfaces at a distance *d* = 26, slightly larger than the contour length of a Ki-67 molecule. The system is initially equilibrated for a time 10^5^ using only the excluded volume interactions and with the two chromosomes held in place. Then, at time *t* = 0, we introduce the electrostatic interaction and simultaneously release the chromosomes, so that they are free to translate along the *x* axis. During the simulation, which lasts a time *t*_tot_ = 6 × 10^5^, the charge of the Ki-67 beads evolves according to Eq. (6). We perform five independent simulation runs (equilibration and production) and average the results over these simulations.

### Simulation of Ki-67 and RNA condensation in solution

To better display how Ki-67 and RNA cluster into condensates in simulations as Ki-67 is dephosphorylated, we perform additional simulations of Ki-67 and RNA molecules in solution. In these simulations, there are no surfaces representing the chromosomes, and thus no grafted Ki-67 molecules. We simulate 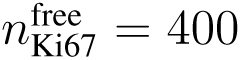 Ki-67 molecules and *n*_RNA_ = 100 RNA molecules.

The charge of Ki-67 is chosen to mimic the fully dephosphorylated state reached in mitotic exit. The total concentration (number of beads per unit volume) is *c* = 10*^−^*^2^, and the simulations are performed in a cubic box (edges *L_x_* = *L_y_* = *L_z_* = 125.1) with periodic boundary conditions in all spatial directions. The system is equilibrated for a time 10^5^ using only the excluded volume interactions before the production run with the full interactions. (duration *t*_tot_ = 2 × 10^5^). For these simulations, we use a slightly larger time step, *δt* = 5 × 10*^−^*^3^ instead of *δt* = 2 × 10*^−^*^3^. As these simulations are performed to qualitatively show the that Ki-67 and RNA cluster into condensates, and given that the simulated system is rather large, we perform a single simulation run.

### Charge prediction for simulations

Ki-67 protein sequence was split into 24 beads of 120 amino acid residues per bead (Supplementary Table 3), omitting the LR-domain. The charge per bead (dephosphorylated) was calculated based on the amino acid sequence using a customized R script based on the net charge calculation of the ‘seqinr’ R package and pK values from EMBOSS considering a pH of 7. To estimate the bead charges in a phosphorylated state, we calculated the fraction of serines and threonine residues (potential phosphorylation sites) of Ki-67ΔLR located in the specific bead sequence and added the charge of the phosphorylation assuming a total of 150 phosphorylation sites and −2 net charge per phosphorylated residue to the charge of the dephosphorylated bead: (−2 × [fraction *S* + *T*] × 150 + bead charge[dephosphorylated]).

### Units conversion

To convert the simulation units into experimental ones, we start by the observation that the brush height during early mitosis, measured as the average root-mean-squared end-to-end distance 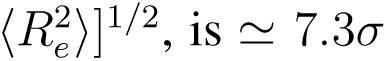 in the simulations. Experimentally, the brush height during early mitosis is approximatley 120 nm. Thus, we set *σ* = 120 nm*/*7.3 = 16 nm. The grafting density of Ki-67 molecules in simulation units is *ρ_g_* = 2.5 10*^−^*^2^*σ^−^*^2^ (for the simulations with the two half cylinders), which corresponds to≃98μm*^−^*^2^ in experimental units. The unit charge 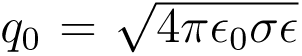 can be estimated by setting *ɛ* (the unit of energy) to be equal to *k_B_T*, with *k_B_* Boltzmann’s constant and *T* = 37*^◦^*C 310 K. In experimental units, *k_B_T* 4.3 10*^−^*^21^ J. The unit charge is thus q_0_ ≃ 8.7 × 10*^−^*^20^ C, which corresponds to 0.55 atomic charge units. In accordance with the unit length conversion, we set the unit mass *m* (mass of a bead of a Ki-67 polymer in the simulations) to be equal to 1/24th of the mass of a Ki-67 molecule, which is 3.58 × 10^5^ Dalton = 5.96 × 10*^−^*^19^ g. Thus, *m* = 2.5 × 10*^−^*^20^ g.

**Table 1:** Units conversions for the main physical quantities. Conversion from MD reduced units (simulation units) to SI units (experimental units).

| Physical quantity | Simulation unit | Experimental unit (SI) |
| --- | --- | --- |
| Length | $\sigma$ | $1.6 \times 10^{-8} \text{ m}$ |
| Charge | $q_0$ | $8.7 \times 10^{-20} \text{ C}$ |
| Energy | $\epsilon$ | $4.3 \times 10^{-21} \text{ J}$ |
| Temperature | $\epsilon/k_B$ | $310 \text{ K}$ |
| Mass | $m$ | $2.5 \times 10^{-20} \text{ g}$ |

---

**Figure S1.**
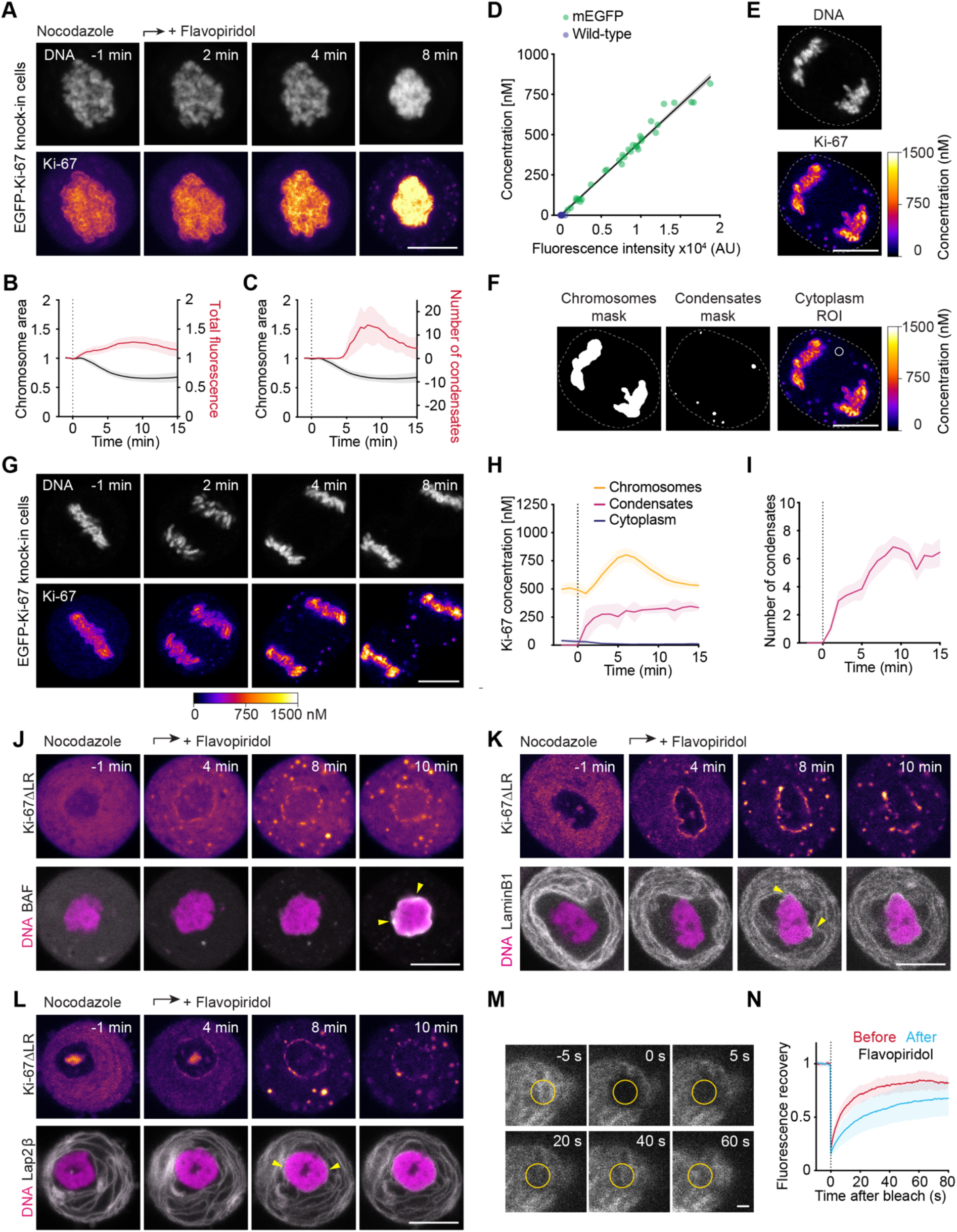
Endogenously tagged Ki-67 phase separates and enriches on chromosomes during mitotic exit prior to nuclear envelope reformation. (A) Time-lapse imaging of endogenously EGFP-tagged Ki-67 during spindle-less mitotic exit. (B) Quantification of chromosome ensemble area and EGFP total fluorescence intensity in segmented chromosome ensemble area over time as in (A). (C) Quantification of endogenously EGFP-tagged Ki-67 cytoplasmic condensates over time as in (A). n = 12. (D–I) Dynamic concentration maps of endogenous Ki-67 during mitotic exit. (D) Calibration line of mEGFP fluorescent intensities and the corresponding protein concentrations as determined by Fluorescence correlation spectroscopy (FCS) measured in nucleoplasm and cytoplasm of untransfected HeLa Kyoto cells or transfected with mEGFP. n = 10 (untransfected wild-type cells); n = 33 (transfected with mEGFP). (E) Fluorescence microscopy image of live HeLa cell undergoing anaphase with endogenously EGFP-labelled Ki-67, scaled to absolute Ki-67 concentration as determined by FCS-calibrated imaging. (F) Example segmentation of chromosomes (left), cytoplasmic condensates (middle) and cytoplasmic ROI (right, white circle) used for quantification in (G–I). (G) EGFP-Ki-67 knock-in cell undergoing anaphase. Images were calibrated by FCS to convert fluorescence intensities into cellular protein concentration (colour bars) as in (D, E). (H) Quantification of EGFP-Ki-67 concentration in chromosomes, cytoplasmic condensates and cytoplasm over anaphase as in (G). n = 13. (I) Quantification of endogenously EGFP-tagged Ki-67 cytoplasmic condensates over time as in (G). (J–L) Time-lapse microscopy of Ki-67 KO cells transfected with EGFP-Ki-67ΔLR and constitutively expressing either BAF-TagRFP (J), TagRFP-LaminB1 (K) or mCherry-Lap2ý (L) during spindle-less mitotic exit. Yellow arrows indicate the earliest BAF, LaminB1 or Lap2ý binding, respectively. (M) Endogenously EGFP-tagged Ki-67 bound to chromosomes was photobleached before and 3 minutes after induction of spindle-less mitotic exit and the recovery of fluorescence was followed by time-lapse recording in a circular image region (yellow circle). Representative example of a cell undergoing mitotic exit quantified in (N). (N) Quantification of EGFP mean fluorescence intensity in the bleached area before flavopiridol (Early mitosis) and after flavopiridol addition (Mitotic exit), normalized to pre-bleached values. Fluorescence recovery measurements: n = 19 (Early mitosis); n = 24 (Mitotic exit). Dashed vertical lines indicate flavopiridol addition (B, C), onset of chromatids segregation (H, I) or photobleaching (N). Lines and shaded areas represent mean ± SD (B-D, H, I, N). DNA was labelled with SiR-Hoechst. Max. intensity z-projections (A) or single z-slices (E–G, J–M) are shown. Scale bars, 10 µm (A, E–G, J–L) or 1 µm (M).

**Figure S2.**
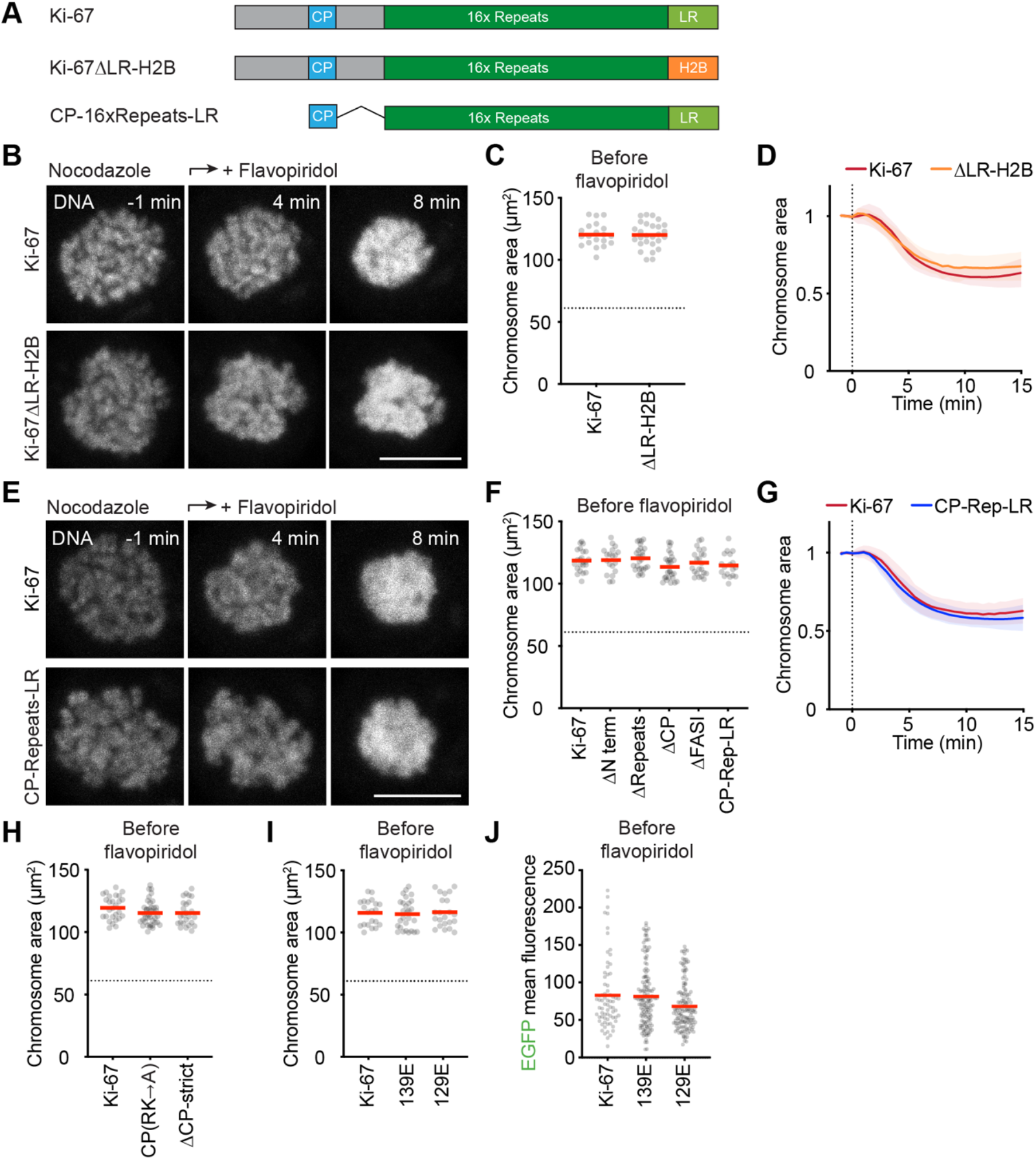
The CP domain is key for chromosome clustering while the LR domain anchors Ki-67 to chromosomes without specifically contributing to chromosome clustering. (A) Schematic of Ki-67 constructs used in (B-G). (B) Time-lapse microscopy images of HeLa Ki-67 KO cells transfected with either full-length EGFP-Ki-67 or EGFP-Ki-67ΔLR-H2B undergoing spindle-less mitotic exit. (C) Initial chromosome ensemble area (prior to flavopiridol addition) of Ki-67 KO cells transfected with either EGFP-Ki-67 or EGFP-Ki-67ΔLR-H2B plotted in (D). (D) Quantification of chromosome ensemble area over time as in (B). n = 20 (Ki-67); n = 27 (Ki-67ΔLR-H2B). (E) Time-lapse microscopy images of HeLa Ki-67 KO cells transfected with either full-length EGFP-Ki-67 or EGFP-CP-Repeats-LR undergoing spindle-less mitotic exit. (F) Initial chromosome ensemble area (prior to flavopiridol addition) of Ki-67 KO cells transfected with EGFP-labelled full-length Ki-67, Ki-67ΔN terminus, Ki-67ΔRepeats, Ki-67ΔCP, Ki-67ΔFASI or CP-Repeats-LR, plotted in Figure 3F and (G). (G) Quantification of chromosome ensemble area over time as in (E). n = 18 (CP-Repeats-LR). (H) Initial chromosome ensemble area (prior to flavopiridol addition) of Ki-67 KO cells transfected with EGFP-labelled full-length Ki-67, Ki-67(CP(RK®A)) or Ki-67ΔCP-strict, plotted in Figure 4G. (I) Initial chromosome ensemble area (prior to flavopiridol addition) of Ki-67 KO cells transfected with EGFP-labelled full-length Ki-67, Ki-67(139E) or Ki-67(129E), plotted in Figure 2E and 5F. (J) Quantification of EGFP mean fluorescence intensity of chromatids prior to flavopiridol addition plotted in Figure 5H. Dashed vertical lines indicate flavopiridol addition (D, G) and dashed horizontal lines dashed horizontal lines indicate mean chromosome ensemble area of Ki-67 KO cells from Figure 1E for reference (C, F, H, I). Red bars represent mean (C, F, H– J). Lines and shaded areas represent mean ± SD (D, G). DNA was labelled with SiR-Hoechst. Max. intensity z-projections are shown. Scale bar, 10 µm (B, E).

**Figure S3.**
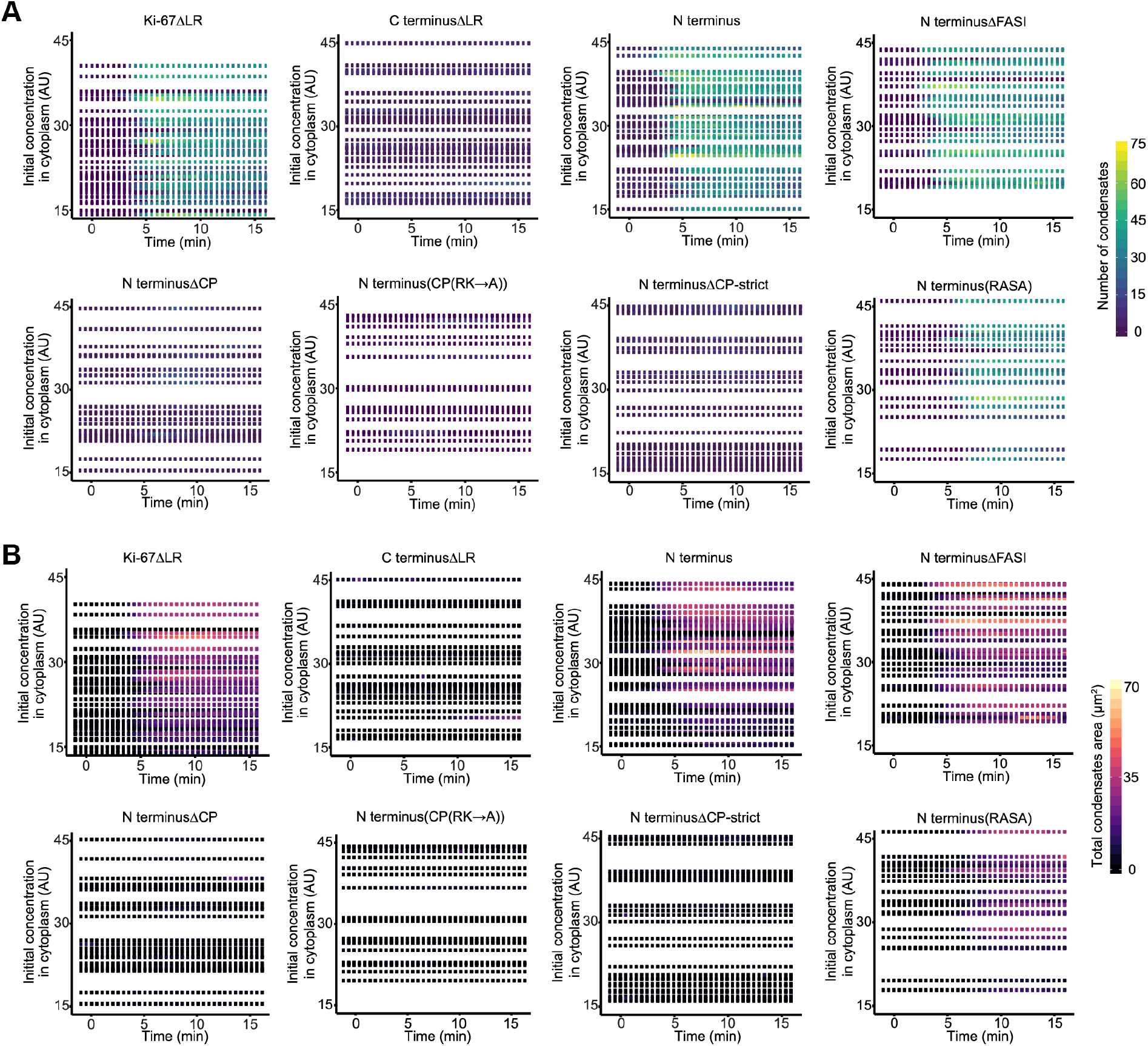
Phase separation dynamics of various Ki-67 mutants relative to protein expression levels. (A, B) Heatmaps displaying number of cytoplasmic condensates (A) or total area of condensates in µm^2^ (B) as a function of time and initial cytoplasmic concentration of cells analysed in Figures 3B, 3D, 4C, 4D and S4C. Time relative to flavopiridol addition.

**Figure S4.**
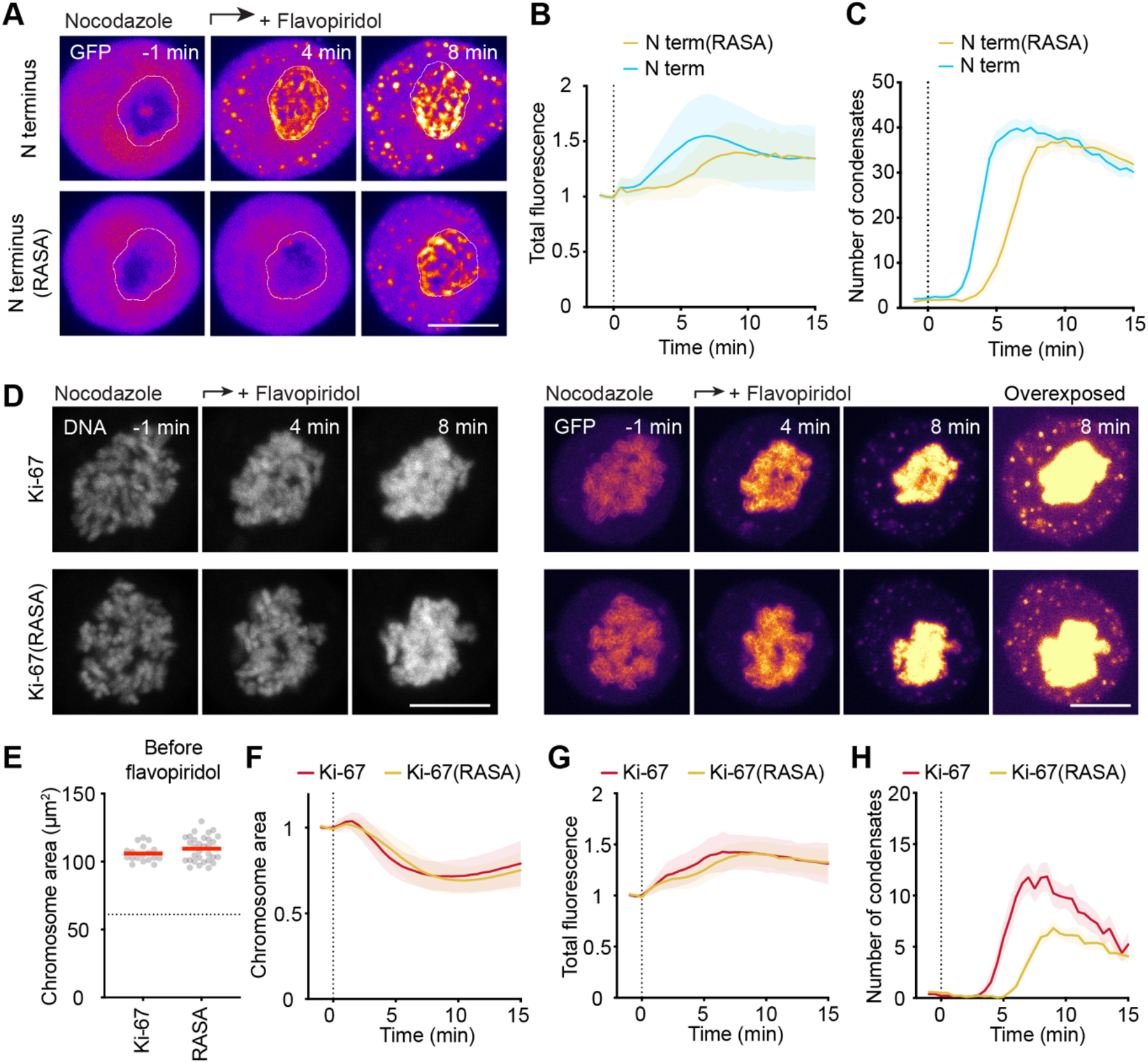
Protein phosphatase binding is not required for chromosome clustering but modulates the timing. (A) Time-lapse microscopy of spindle-less mitotic exit of HeLa Ki-67 KO cell transiently expressing Ki-67 N terminus or N terminus(RASA) constructs tagged with EGFP. White lines represent chromosomal regions used for quantifications in (B). (B) Quantification of EGFP total fluorescence intensity in segmented chromosome ensemble area over time as in (A). (C) Quantification of number of cytoplasmic condensates over time as in (A). (N terminus for reference as in Figure 3C and 3D); n = 22 (N terminus(RASA)). (D) Time-lapse microscopy images of HeLa EGFP-Ki-67 knock-in cells (top) or HeLa EGFP-Ki-67(RASA) knock-in cells (bottom) undergoing spindle-less mitotic exit. DNA was labelled with SiR-Hoechst. (E) Initial chromosome ensemble area (prior to flavopiridol addition) of HeLa EGFP-Ki-67 knock-in cells or HeLa EGFP-Ki-67(RASA) knock-in cells plotted in (F). Red bars represents mean. Dashed horizontal line indicates mean chromosome ensemble area of Ki-67 KO cells from Figure 1E for reference. (F) Quantification of chromosome ensemble area over time as in (D). (G) Quantification of EGFP total fluorescence intensity in chromosome ensemble area over time as in (D). (H) Quantification of number of cytoplasmic condensates over time as in (D). n = 18 (Ki-67); n = 31 (Ki-67(RASA)). Dashed vertical lines indicate flavopiridol addition (B, C, F, G, H). Lines and shaded areas represent mean ± SD (B, F, G) or mean ± SEM (C, H). Max. intensity z-projections are shown. Scale bars, 10 µm (A, D).

**Figure S5.**
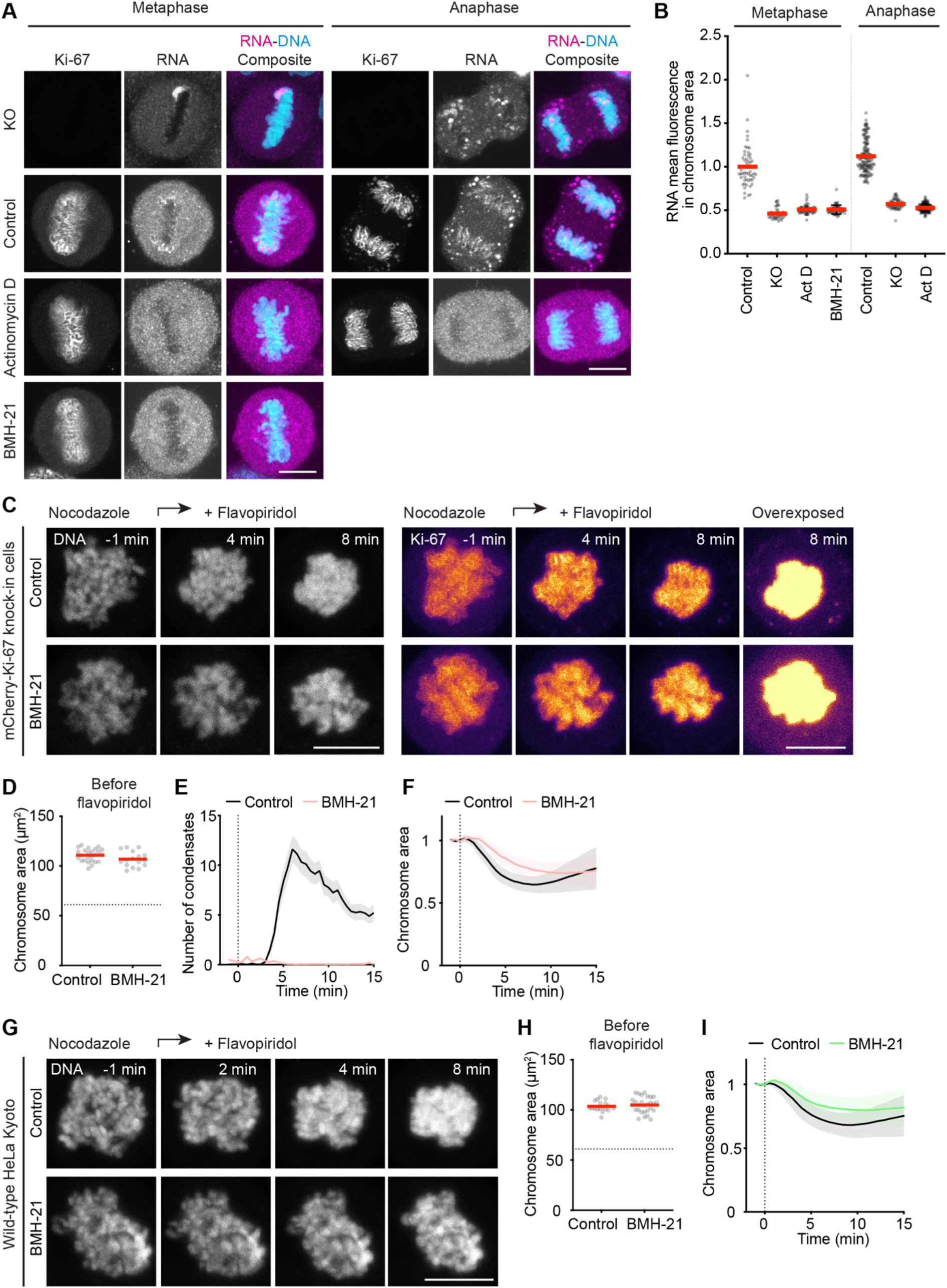
Depletion of pre-rRNA neither affects chromosome individualization nor Ki-67 enrichment on anaphase chromosomes, but impairs Ki-67 phase separation and chromosome clustering. (A) Representative 5’ETS pre-rRNA-FISH images of Ki-67 KO cells or EGFP-Ki-67 knock-in cells untreated (Control) or treated with either actinomycin D (Act D) or BMH-21 undergoing metaphase or anaphase. Notice that BMH-21 treated cells failed to undergo anaphase. (B) Quantification of RNA mean fluorescence intensity in segmented chromosome area as in (A). Values were normalized to median value of control cells undergoing metaphase. Metaphase: n = 53 (Control); n = 39 (KO); n = 68 (Act D); n = 51 (BMH-21). Anaphase: n = 140 (Control); n = 55 (KO); n = 145 (Act D). (C) Time-lapse images of spindle-less mitotic exit of mCherry-Ki-67 knock-in cells. Cells were untreated (Control) or treated with BMH-21. (D) Initial chromosome ensemble area (prior to flavopiridol addition) of HeLa mCherry-Ki-67 knock-in cells plotted in (F). (E) Quantification of number of cytoplasmic condensates over time as in (C). (F) Quantification of chromosome ensemble area over time as in (C). n = 25 (Control); n = 16 (BMH-21). (G) Time-lapse images of spindle-less mitotic exit Wild-type HeLa Kyoto cells. Cells were untreated (Control) or treated with BMH-21. (H) Initial chromosome ensemble area (prior to flavopiridol addition) of Wild-type HeLa Kyoto cells plotted in (I). (I) Quantification of chromosome ensemble area over time as in (G). n = 18 (Control); n = 28 (BMH-21). Dashed vertical lines indicate flavopiridol addition (E, F, I) and dashed horizontal lines indicate mean chromosome ensemble area of Ki-67 KO cells from Figure 1E for reference (D, H). Red bars indicate mean (B, D, H). Lines and shaded areas represent mean ± SEM (E) or mean ± SD (F, I). DNA was labelled with DAPI (A) or SiR-Hoechst (C, G). Max. intensity z-projections are shown. Scale bars, 10 µm (A, C, G).

**Figure S6.**
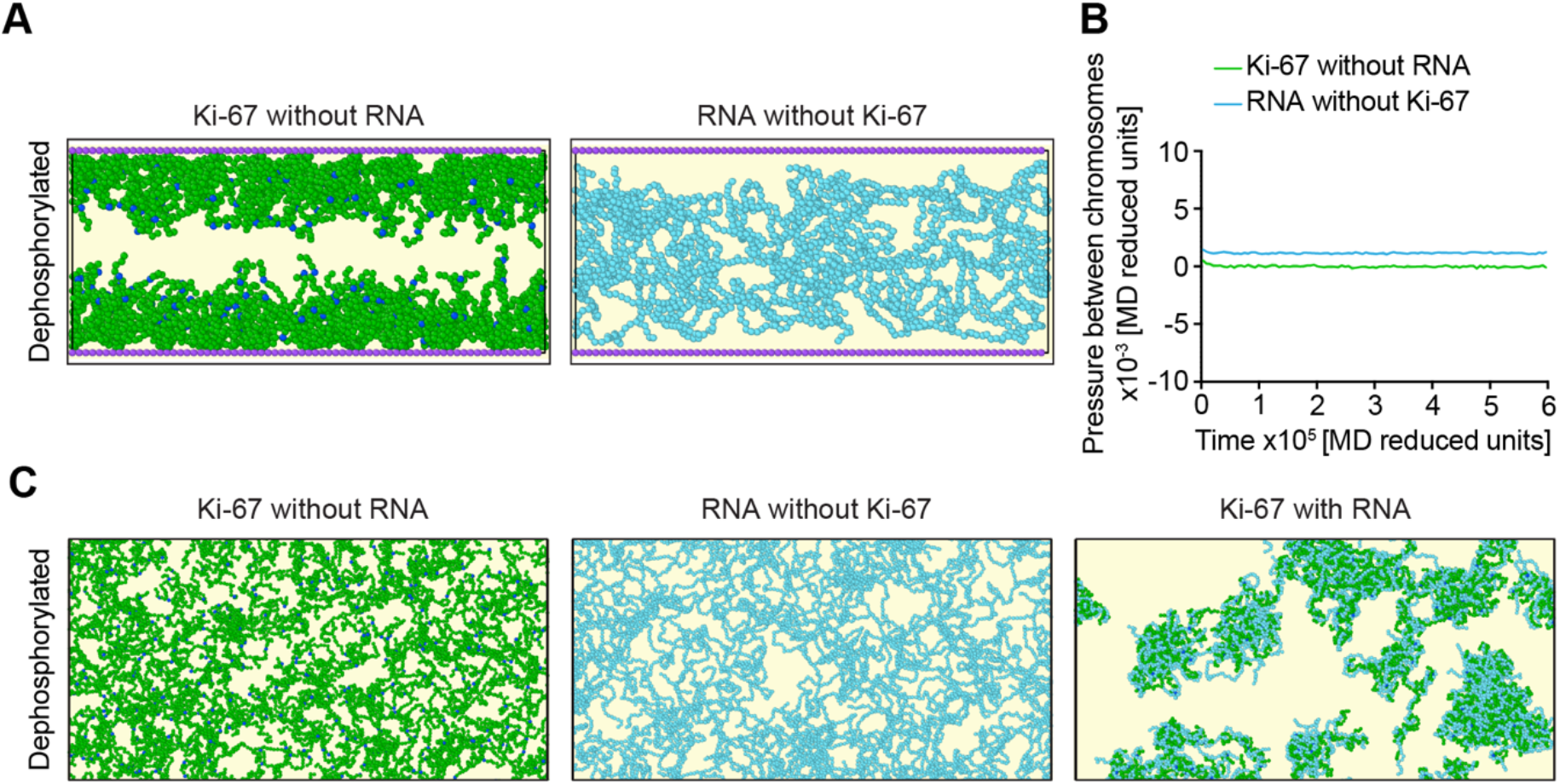
RNA and dephosphorylated Ki-67 are required for chromosome attraction and condensate formation in coarse-grained simulations. (A, B) Simulations of forces acting between two chromosome surfaces as the charge of Ki-67 is gradually increased as in Figure 6A. (A) The two chromosomes are modelled as two immobile surfaces (violet), with a constant slightly negative charge, decorated with bead-spring polymers representing Ki-67 molecules (green – CP in dark blue) and constant highly negatively charged RNA molecules (light blue). Simulation snapshots for dephosphorylated Ki-67 or RNA alone. (B) Pressure (force per unit area) between chromosomes as a function of simulation time using the gradual Ki-67 charge increase as in (Figure /A). Lines and shaded areas represent mean ± SD. n = 5 (Ki-67 without RNA); n = 5 (RNA without Ki-67). (C) Simulations of Ki-67 and RNA condensation in solution as the charge of Ki-67 is gradually increased as in Figure 6A. Simulation snapshots of dephosphorylated Ki-67 (green – CP in dark blue) with or without RNA (light blue), in addition to RNA without Ki-67.

**Figure S7.**
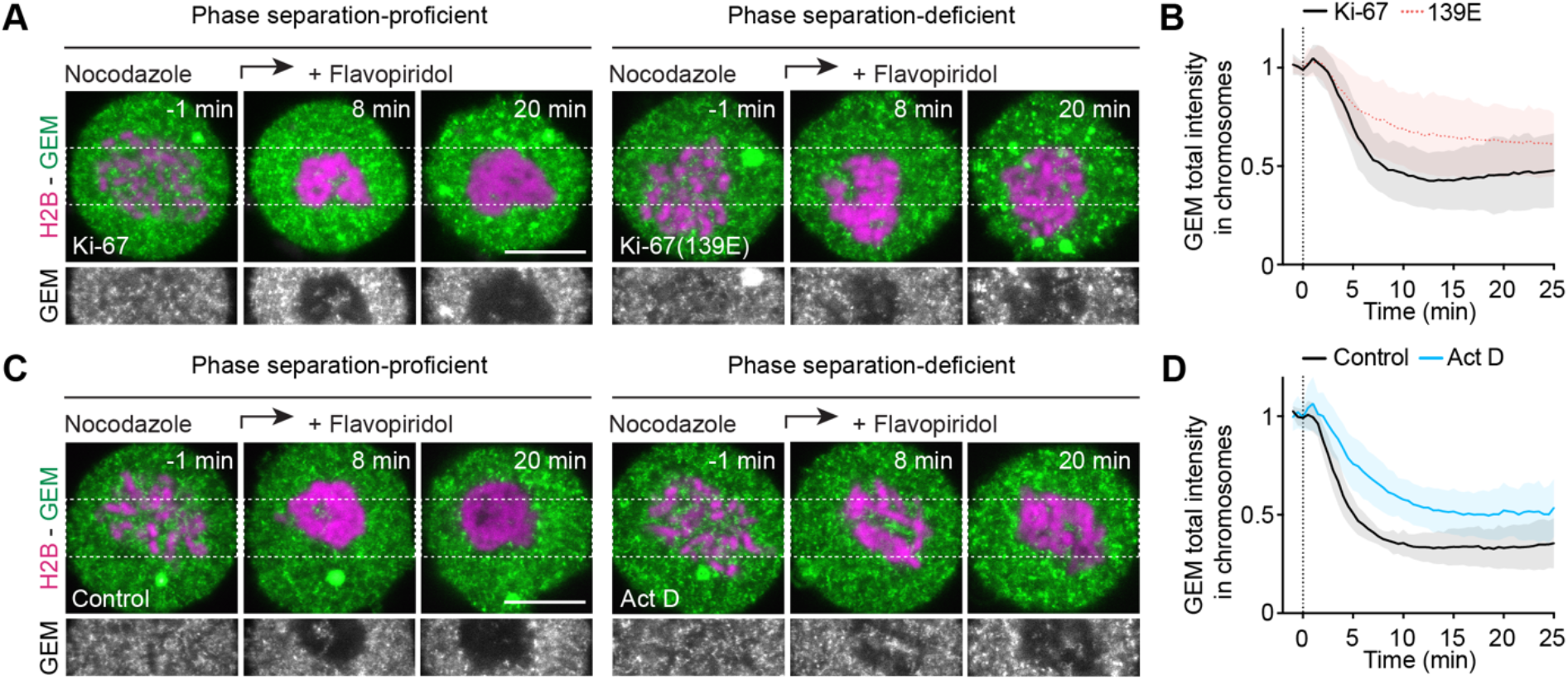
Preventing Ki-67 phase separation by phosphomimetic mutations or rRNA depletion prevents GEM exclusion from the future nuclear space. (A) Time-lapse microscopy of spindle-less mitotic exit in Ki-67 KO cells stably expressing GEMs and transiently expressing Ki-67 full-length (left) or Ki-67(139E) (right). (B) Quantification of GEM exclusion over time as in (A). n = 25 (Ki-67); n = 28 (Ki-67(139E)). (C) Time-lapse microscopy of spindle-less mitotic exit in Ki-67 wild-type cells stably expressing GEMs under Control (left) or actinomycin D (Act D) treatment (right) conditions. (D) Quantification of GEM exclusion over time as in (C). n = 27 (Control); n = 31 (Act D). Images show timepoint 1 min before and after 8 min and 20 min of flavopiridol-induced mitotic exit. Dashed vertical lines indicate flavopiridol addition (B, D). Lines and shaded areas represent mean ± SD (B, D). Single z-slices are shown. Scale bars, 10 µm (A, C).

## References

1. Paulson, J.R., Hudson, D.F., Cisneros-Soberanis, F., and Earnshaw, W.C. (2021). Mitotic chromosomes. Semin Cell Dev Biol 117, 7–29. 10.1016/j.semcdb.2021.03.014.

2. Zanden, S.Y. van der, Jongsma, M.L.M., Neefjes, A.C.M., Berlin, I., and Neefjes, J. (2022). Maintaining soluble protein homeostasis between nuclear and cytoplasmic compartments across mitosis. Trends Cell Biol. 10.1016/j.tcb.2022.06.002.

3. Booth, D.G., and Earnshaw, W.C. (2017). Ki-67 and the Chromosome Periphery Compartment in Mitosis. Trends Cell Biol 27, 906–916. 10.1016/j.tcb.2017.08.001.

4. Stenström, L., Mahdessian, D., Gnann, C., Cesnik, A.J., Ouyang, W., Leonetti, M.D., Uhlén, M., Cuylen-Haering, S., Thul, P.J., and Lundberg, E. (2020). Mapping the nucleolar proteome reveals a spatiotemporal organization related to intrinsic protein disorder. Mol Syst Biol 16, e9469. 10.15252/msb.20209469.

5. Hernandez-Verdun, D. le, and Gautier, T. (1994). The chromosome periphery during mitosis. Bioessays 16, 179–185. 10.1002/bies.950160308.

6. Booth, D.G., Takagi, M., Sanchez-Pulido, L., Petfalski, E., Vargiu, G., Samejima, K., Imamoto, N., Ponting, C.P., Tollervey, D., Earnshaw, W.C., et al. (2014). Ki-67 is a PP1-interacting protein that organises the mitotic chromosome periphery. Elife 3, e01641.

7. Hayashi, Y., Kato, K., and Kimura, K. (2017). The hierarchical structure of the perichromosomal layer comprises Ki67, ribosomal RNAs, and nucleolar proteins. Biochem Biophys Res Commun 493, 1043–1049. 10.1016/j.bbrc.2017.09.092.

8. Cuylen, S., Blaukopf, C., Politi, A.Z., Müller-Reichert, T., Neumann, B., Poser, I., Ellenberg, J., Hyman, A.A., and Gerlich, D.W. (2016). Ki-67 acts as a biological surfactant to disperse mitotic chromosomes. Nature 535, 308–312. 10.1038/nature18610.

9. Verheijen, R., Kuijpers, H.J., Driel, R. van, Beck, J.L., Dierendonck, J.H. van, Brakenhoff, G.J., and Ramaekers, F.C. (1989). Ki-67 detects a nuclear matrix-associated proliferation-related antigen. II. Localization in mitotic cells and association with chromosomes. J Cell Sci 92, 531–540.

10. Uhlmann, F., Lottspeich, F., and Nasmyth, K. (1999). Sister-chromatid separation at anaphase onset is promoted by cleavage of the cohesin subunit Scc1. Nature 400, 37–42. 10.1038/21831.

11. Cuylen-Haering, S., Petrovic, M., Hernandez-Armendariz, A., Schneider, M.W.G., Samwer, M., Blaukopf, C., Holt, L.J., and Gerlich, D.W. (2020). Chromosome clustering by Ki-67 excludes cytoplasm during nuclear assembly. Nature 587, 285–290. 10.1038/s41586-020-2672-3.

12. Banani, S.F., Lee, H.O., Hyman, A.A., and Rosen, M.K. (2017). Biomolecular condensates: organizers of cellular biochemistry. Nat Rev Mol Cell Biol 18, 285–298. 10.1038/nrm.2017.7.

13. Yamazaki, H., Takagi, M., Kosako, H., Hirano, T., and Yoshimura, S.H. (2022). Cell cycle-specific phase separation regulated by protein charge blockiness. Nat Cell Biol 24, 625– 632. 10.1038/s41556-022-00903-1.

14. Endl, E., and Gerdes, J. (2000). Posttranslational modifications of the KI-67 protein coincide with two major checkpoints during mitosis. J. Cell. Physiol. 182, 371–380. 10.1002/(sici)1097-4652(200003)182:3<371::aid-jcp8>3.0.co;2-j.

15. Potapova, T.A., Daum, J.R., Pittman, B.D., Hudson, J.R., Jones, T.N., Satinover, D.L., Stukenberg, P.T., and Gorbsky, G.J. (2006). The reversibility of mitotic exit in vertebrate cells. Nature 440, 954–958. 10.1038/nature04652.

16. Takagi, M., Matsuoka, Y., Kurihara, T., and Yoneda, Y. (1999). Chmadrin: a novel Ki-67 antigen-related perichromosomal protein possibly implicated in higher order chromatin structure. J Cell Sci 112 ( *Pt 15*), 2463–2472.

17. Samwer, M., Schneider, M.W.G., Hoefler, R., Schmalhorst, P.S., Jude, J.G., Zuber, J., and Gerlich, D.W. (2017). DNA Cross-Bridging Shapes a Single Nucleus from a Set of Mitotic Chromosomes. Cell 170, 956–972.e23. 10.1016/j.cell.2017.07.038.

18. Saiwaki, T., Kotera, I., Sasaki, M., Takagi, M., and Yoneda, Y. (2005). In vivo dynamics and kinetics of pKi-67: Transition from a mobile to an immobile form at the onset of anaphase. Exp Cell Res 308, 123–134. 10.1016/j.yexcr.2005.04.010.

19. Dephoure, N., Zhou, C., Villén, J., Beausoleil, S.A., Bakalarski, C.E., Elledge, S.J., and Gygi, S.P. (2008). A quantitative atlas of mitotic phosphorylation. Proc Natl Acad Sci USA 105, 10762–10767. 10.1073/pnas.0805139105.

20. Takagi, M., Nishiyama, Y., Taguchi, A., and Imamoto, N. (2014). Ki67 Antigen Contributes to the Timely Accumulation of Protein Phosphatase 1γ on Anaphase Chromosomes*. J Biol Chem 289, 22877–22887. 10.1074/jbc.m114.556647.

21. Wootton, J.C. (1994). Non-globular domains in protein sequences: automated segmentation using complexity measures. Comput. Chem. 18, 269–285.

22. Schlüter, C., Duchrow, M., Wohlenberg, C., Becker, M.H., Key, G., Flad, H.D., and Gerdes, J. (1993). The cell proliferation-associated antigen of antibody Ki-67: a very large, ubiquitous nuclear protein with numerous repeated elements, representing a new kind of cell cycle-maintaining proteins. J Cell Biol 123, 513–522.

23. Roden, C., and Gladfelter, A.S. (2021). RNA contributions to the form and function of biomolecular condensates. Nat Rev Mol Cell Biol 22, 183–195. 10.1038/s41580-020-0264-6.

24. Cochard, A., Navarro, M.G.-J., Piroska, L., Kashida, S., Kress, M., Weil, D., and Gueroui, Z. (2022). RNA at the surface of phase-separated condensates impacts their size and number. Biophys J 121, 1675–1690. 10.1016/j.bpj.2022.03.032.

25. Gautier, T., Robert-Nicoud, M., Guilly, M.N., and Hernandez-Verdun, D. (1992). Relocation of nucleolar proteins around chromosomes at mitosis. A study by confocal laser scanning microscopy. J Cell Sci 102 ( *Pt 4*), 729–737.

26. Ma, K., Luo, M., Xie, G., Wang, X., Li, Q., Gao, L., Yu, H., and Yu, X. (2022). Ribosomal RNA regulates chromosome clustering during mitosis. Cell Discov 8, 51. 10.1038/s41421-022-00400-7.

27. Takagi, M., Sueishi, M., Saiwaki, T., Kametaka, A., and Yoneda, Y. (2001). A novel nucleolar protein, NIFK, interacts with the forkhead associated domain of Ki-67 antigen in mitosis. J Biol Chem 276, 25386–25391. 10.1074/jbc.m102227200.

28. Hooser, A.A.V., Yuh, P., and Heald, R. (2005). The perichromosomal layer. Chromosoma 114, 377–388. 10.1007/s00412-005-0021-9.

29. Booth, D.G., Beckett, A.J., Molina, O., Samejima, I., Masumoto, H., Kouprina, N., Larionov, V., Prior, I.A., and Earnshaw, W.C. (2016). 3D-CLEM Reveals that a Major Portion of Mitotic Chromosomes Is Not Chromatin. Mol Cell 64, 790–802. 10.1016/j.molcel.2016.10.009.

30. Pak, C.W., Kosno, M., Holehouse, A.S., Padrick, S.B., Mittal, A., Ali, R., Yunus, A.A., Liu, D.R., Pappu, R.V., and Rosen, M.K. (2016). Sequence Determinants of Intracellular Phase Separation by Complex Coacervation of a Disordered Protein. Mol. Cell 63, 72–85. 10.1016/j.molcel.2016.05.042.

31. Boeynaems, S., Holehouse, A.S., Weinhardt, V., Kovacs, D., Lindt, J.V., Larabell, C., Bosch, L.V.D., Das, R., Tompa, P.S., Pappu, R.V., et al. (2019). Spontaneous driving forces give rise to protein−RNA condensates with coexisting phases and complex material properties. Proc. Natl. Acad. Sci. 116, 7889–7898. 10.1073/pnas.1821038116.

32. Quail, T., Golfier, S., Elsner, M., Ishihara, K., Murugesan, V., Renger, R., Jülicher, F., and Brugués, J. (2021). Force generation by protein–DNA co-condensation. Nat Phys 17, 1007–1012. 10.1038/s41567-021-01285-1.

33. Keenen, M.M., Brown, D., Brennan, L.D., Renger, R., Khoo, H., Carlson, C.R., Huang, B., Grill, S.W., Narlikar, G.J., and Redding, S. (2021). HP1 proteins compact DNA into mechanically and positionally stable phase separated domains. Elife 10, e64563. 10.7554/elife.64563.

34. Renger, R., Morin, J.A., Lemaitre, R., Ruer-Gruss, M., Jülicher, F., Hermann, A., and Grill, S.W. (2022). Co-condensation of proteins with single- and double-stranded DNA. Proc National Acad Sci 119, e2107871119. 10.1073/pnas.2107871119.

35. Shin, Y., Chang, Y.-C., Lee, D.S.W., Berry, J., Sanders, D.W., Ronceray, P., Wingreen, N.S., Haataja, M., and Brangwynne, C.P. (2018). Liquid Nuclear Condensates Mechanically Sense and Restructure the Genome. Cell 175, 1481–1491.e13. 10.1016/j.cell.2018.10.057.

36. Gennes, P.-G. de, Brochard-Wyart, F., and Quéré, D. (2004). Capillarity and Wetting Phenomena, Drops, Bubbles, Pearls, Waves. 1–31. 10.1007/978-0-387-21656-0_1.

37. Steinberg, M.S. (1996). Adhesion in Development: An Historical Overview. Dev Biol 180, 377–388. 10.1006/dbio.1996.0312.

38. Hayashi, T., and Carthew, R.W. (2004). Surface mechanics mediate pattern formation in the developing retina. Nature 431, 647–652. 10.1038/nature02952.

39. Gouveia, B., Kim, Y., Shaevitz, J.W., Petry, S., Stone, H.A., and Brangwynne, C.P. (2022). Capillary forces generated by biomolecular condensates. Nature 609, 255–264. 10.1038/s41586-022-05138-6.

40. Broyles, B.K., Gutierrez, A.T., Maris, T.P., Coil, D.A., Wagner, T.M., Wang, X., Kihara, D., Class, C.A., and Erkine, A.M. (2021). Activation of gene expression by detergent-like protein domains. Iscience 24, 103017. 10.1016/j.isci.2021.103017.

41. Kelley, F.M., Favetta, B., Regy, R.M., Mittal, J., and Schuster, B.S. (2021). Amphiphilic proteins coassemble into multiphasic condensates and act as biomolecular surfactants. Proc National Acad Sci 118, e2109967118. 10.1073/pnas.2109967118.

42. Folkmann, A.W., Putnam, A., Lee, C.F., and Seydoux, G. (2021). Regulation of biomolecular condensates by interfacial protein clusters. Science 373, 1218–1224. 10.1126/science.abg7071.

43. Schmitz, M.H.A., Held, M., Janssens, V., Hutchins, J.R.A., Hudecz, O., Ivanova, E., Goris, J., Trinkle-Mulcahy, L., Lamond, A.I., Poser, I., et al. (2010). Live-cell imaging RNAi screen identifies PP2A-B55[alpha] and importin-[beta]1 as key mitotic exit regulators in human cells. Nat Cell Biol 12, 886–893. 10.1038/ncb2092.

44. Lukinavičius, G., Blaukopf, C., Pershagen, E., Schena, A., Reymond, L., Derivery, E., Gonzalez-Gaitan, M., D’Este, E., Hell, S.W., Gerlich, D.W., et al. (2015). SiR–Hoechst is a far-red DNA stain for live-cell nanoscopy. Nat Comms 6, 8497. 10.1038/ncomms9497.

45. Carron, C., O’Donohue, M.-F., Choesmel, V., Faubladier, M., and Gleizes, P.-E. (2011). Analysis of two human pre-ribosomal factors, bystin and hTsr1, highlights differences in evolution of ribosome biogenesis between yeast and mammals. Nucleic Acids Res 39, 280–291. 10.1093/nar/gkq734.

46. Sommer, C., Hoefler, R., Samwer, M., and Gerlich, D.W. (2017). A deep learning and novelty detection framework for rapid phenotyping in high-content screening. Mol Biol Cell 28, 3428–3436. 10.1091/mbc.e17-05-0333.

47. Politi, A.Z., Cai, Y., Walther, N., Hossain, M.J., Koch, B., Wachsmuth, M., and Ellenberg, J. (2018). Quantitative mapping of fluorescently tagged cellular proteins using FCS-calibrated four-dimensional imaging. Nat. Protoc. 13, 1445–1464. 10.1038/nprot.2018.040.

48. Berg, S., Kutra, D., Kroeger, T., Straehle, C.N., Kausler, B.X., Haubold, C., Schiegg, M., Ales, J., Beier, T., Rudy, M., et al. (2019). ilastik: interactive machine learning for (bio)image analysis. Nat. Methods 16, 1226–1232. 10.1038/s41592-019-0582-9.

49. Plimpton, S. (1995). Fast Parallel Algorithms for Short-Range Molecular Dynamics. J Comput Phys 117, 1–19. 10.1006/jcph.1995.1039.

50. Kremer, K., and Grest, G.S. (1990). Dynamics of entangled linear polymer melts: A molecular-dynamics simulation. J Chem Phys 92, 5057–5086. 10.1063/1.458541.

51. Svaneborg, C., and Everaers, R. (2020). Characteristic Time and Length Scales in Melts of Kremer–Grest Bead–Spring Polymers with Wormlike Bending Stiffness. Macromolecules 53, 1917–1941. 10.1021/acs.macromol.9b02437.

52. Schneider, T., and Stoll, E. (1978). Molecular-dynamics study of a three-dimensional one-component model for distortive phase transitions. Phys Rev B 17, 1302–1322. 10.1103/physrevb.17.1302.

