## Supplemental Information for "A liquid-like coat mediates chromosome clustering during mitotic exit"

### Supplemental Video legends

**Video S1. Ki-67 and Ki-67ΔLR enrich on chromosomes and phase separate during mitotic exit.** Time-lapse microscopy of nocodazole arrested cells undergoing spindle-less mitotic exit upon addition of flavopiridol (0 min). Ki-67 KO cells transiently expressing either full-length EGFP-Ki-67 (left) or EGFP-Ki-67ΔLR (right). Maximum intensity z-projections of EGFP (top) and DNA stained with SiR-Hoechst (bottom) are shown. Scale bars, 10 μm. Same data shown in Figure 1.

**Video S2. Cytoplasmic Ki-67ΔLR foci fuse with the condensed Ki-67ΔLR phase on the chromosome surface.** Time-lapse microscopy of Ki-67 wild-type cells overexpressing Ki-67ΔLR-mNeonGreen. A single z-slice is shown. Time relative to anaphase onset. Scale bar, 10 μm. Same data shown in Figure 1.

**Video S3. Phosphomimetic Ki-67 fails to phase separate and cluster chromosomes during mitotic exit.** Time-lapse microscopy of nocodazole arrested cells undergoing spindle-less mitotic exit upon addition of flavopiridol (0 min). Ki-67 KO cells transiently expressing either full-length Ki-67 (left) or Ki-67(139E) (right) tagged with EGFP. Maximum intensity z-projections of DNA stained with SiR-Hoechst (top) and EGFP (bottom) are shown. Scale bars, 10 μm. Same data shown in Figure 2.

**Video S4. Phosphomimetic Ki-67ΔLR fails to phase separate and enrich on chromosomes during mitotic exit.** Time-lapse microscopy of nocodazole arrested cells undergoing spindle-less mitotic exit upon addition of flavopiridol (0 min). Ki-67 KO cells transiently expressing either Ki-67ΔLR (left) or Ki-67(139E)ΔLR (right) tagged with EGFP. Maximum intensity z-projections of EGFP (top) and DNA stained with SiR-Hoechst (bottom) are shown. Scale bars, 10 μm. Same data shown in Figure 2.

**Video S5. Phase separation of Ki-67 depends on a positively charged patch within its N terminus.** Time-lapse microscopy of nocodazole arrested cells undergoing spindle-less mitotic exit upon addition of flavopiridol (0 min). Ki-67 KO cells transiently expressing either the N-terminal segment (left), the N-terminal segment lacking the charged patch (N-termΔCP) (middle) or the N-terminal segment with arginine and lysine residues within its CP substituted to alanine residues (N-term(CP(RK→A))) (right). Maximum intensity z-projections of EGFP (top) and DNA stained with SiR-Hochest (bottom) are shown. Scale bars, 10 μm. Same data shown in Figure 3 and 4.

**Video S6. The positively charged patch within Ki-67 N terminus is required for chromosome clustering during mitotic exit.** Time-lapse microscopy of nocodazole arrested cells undergoing spindle-less mitotic exit upon addition of flavopiridol (0 min). Ki-67 KO cells transiently expressing either full-length Ki-67 (left), Ki-67 lacking the charged patch (Ki-67ΔCP) (middle) or Ki-67 with arginine and lysine residues within its CP substituted to alanine residues (Ki-67(CP(RK→A))) (right). Maximum intensity z-projections of DNA stained with SiR-Hoechst (top) and EGFP (bottom) are shown. Scale bars, 10 µm. Same data shown in Figure 3 and 4.

**Video S7. Depletion of ribosomal RNA reduces Ki-67 phase separation and chromosome clustering.** Time-lapse microscopy of nocodazole arrested EGFP-Ki-67 knock-in cells undergoing spindle-less mitotic exit upon addition of flavopiridol (0 min) under control (left) and actinomycin-D [5 nM] treatment (right) conditions. Maximum intensity z-projections of EGFP (top) and DNA stained with SiR-Hoechst (bottom) are shown. Scale bars, 10 µm. Same data shown in Figure 6.

**Video S8. Chromosome interactions switch from repulsive to attractive in simulations as Ki-67's charge increases.** Simulation movie (top view) of the two chromosomes, modeled as half cylinders (violet) interacting through Ki-67 brushes and RNA. RNA molecules (light blue) are shown as semi-transparent for clarity. The color of Ki-67 molecules reflects their charge (red = negative, grey = neutral, green = positive). The color of the CP (dark blue) is left unchanged for clarity. Same data shown in Figure 7.

**Video S9. Interaction between chromosomes remains repulsive in simulations in the absence of RNA.** Simulation movie (top view) of the two chromosomes, modeled as half cylinders (violet) interacting through Ki-67 brushes. In this simulation, RNA is absent. The color of Ki-67 molecules reflects their charge (red = negative, grey = neutral, green = positive). The color of the CP (dark blue) is left unchanged for clarity. Same data shown in Figure 7.

**Supplemental Table 1. HeLa Kyoto cell lines used in this study.**

| Cell line name | Comments | Reference | Used in Figures | Lab ID |
|---|---|---|---|---|
| Ki-67 KO | Ki-67 knock-out | Published in[8] | Figures 1–6 and S1–S5 | 63 |
| Wild-type | Wild-type cells | Originally from S. Narumiya (Kyoto University, Japan), validated by a Multiplex human Cell line Authentication test (MCA), 21.04.2016 | Figure 1 | 1 |
| EGFP-MKI67 (homozygous) | Endogenously homozygously labelled Ki-67 with N-terminal EGFP | Published in[8] | Figures 6, S1, S4 and S5 | 77 |
| Ki-67 KO (RIEP), BAF-TagRFP | Ki-67 knock-out overexpressing BAF-TagRFP | Gift from Gerlich lab (IMBA, Austria) | Figure S1 | 116 (1662) |
| Ki-67 KO (RIEP), mCherry-Lap2beta | Ki-67 knock-out overexpressing mCherry-Lap2beta | Gift from Gerlich lab (IMBA, Austria) | Figure S1 | 115 (1661) |
| Ki-67 KO (RIEP), TagRFP-LaminB1 | Ki-67 knock-out overexpressing TagRFP-LaminB1 | Gift from Gerlich lab (IMBA, Austria) | Figure S1 | 117 (1663) |
| EGFP-MKI67(RASA) (homozygous) | Endogenously and homozygously labelled Ki-67(RASA) with EGFP at its N terminus | Gift from Gerlich lab (IMBA, Austria) | Figure S4 | 102 (1600) |
| mCherry-MKI67 (homozygous) | Endogenously and homozygously labelled Ki-67 with N-terminal mCherry | Generated in this study | Figure S5 | 149 |
| Ki-67 KO, GEM-EGFP, H2B-mCherry | Ki-67 knock-out expressing genetically encoded multimeric nanoparticles (GEM) and low expression of H2B-mCherry | Gift from Gerlich lab (IMBA, Austria) | Figure S7 | 171 (1698) |
| GEM-EGFP, H2B-mCherry (RIEP) | HeLa expressing GEM-EGFP and low expression of H2B-mCherry | Gift from Gerlich lab (IMBA, Austria) | Figure S7 | 169 (1701) |

**Supplemental Table 2. Plasmids used in this study.**

| Name | Reference | Used in Figures | Lab ID |
|---|---|---|---|
| EGFP-Ki-67 | Generated in this study | Figures 1–5 and S2 | 343 |
| EGFP-Ki-67ΔLR | Generated in this study | Figures 1–3, 5, S1 and S3 | 428 |
| EGFP-LR | Generated in this study | Figure 1 | 726 |
| Ki-67ΔLR-mNeonGreen | Published in[8] | Figure 1 | 190 |
| EGFP-Ki-67(139E) | Generated in this study | Figures 2, 5 and S2 | 318 |
| EGFP-Ki-67(139E)ΔLR | Generated in this study | Figures 2 and 5 | 429 |
| EGFP-N term | Generated in this study | Figures 3, 4, S3 and S4 | 435 |
| EGFP-Ki-67ΔN term | Generated in this study | Figures 3 and S2 | 665 |
| EGFP-Ki-67ΔRepeats | Generated in this study | Figures 3 and S2 | 607 |
| EGFP-Ki-67ΔCP | Generated in this study | Figures 3 and S2 | 483 |
| EGFP-Ki-67ΔFASI | Generated in this study | Figures 3 and S2 | 666 |
| EGFP-Ki-67(C termΔLR) | Generated in this study | Figures 3 and S3 | 434 |
| EGFP-N termΔCP | Generated in this study | Figures 3 and S3 | 475 |
| EGFP-N termΔFASI | Generated in this study | Figures 3 and S3 | 439 |
| EGFP-Ki-67(CP(RK→A)) | Generated in this study | Figures 4 and S2 | 744 |
| EGFP-Ki-67ΔCP-strict | Generated in this study | Figures 4 and S2 | 490 |
| EGFP-N term(CP(RK→A)) | Generated in this study | Figures 4 and S3 | 986 |
| EGFP-N termΔCP-strict | Generated in this study | Figures 4 and S3 | 484 |
| EGFP-Ki-67(129E) | Generated in this study | Figures 5 and S2 | 537 |
| EGFP-Ki-67(129E)ΔLR | Generated in this study | Figures 5 and S2 | 991 |
| mCherry-Ki-67-EGFP | Published in[8] | Figures 5 and S2 | 216 |
| mCherry-Ki-67(139E)-EGFP | Generated in this study | Figures 5 and S2 | 825 |
| mCherry-Ki-67(129E)-EGFP | Generated in this study | Figures 5 and S2 | 922 |
| EGFP-Ki-67ΔLR-H2B | Generated in this study | Figure S2 | 711 |
| EGFP-CP-Repeats-LR | Generated in this study | Figure S2 | 818 |
| EGFP-N term(RASA) | Generated in this study | Figures S3 and S4 | 936 |
| mCherry-Ki-67 | Generated in this study | Figure S7 | 339 |
| mCherry-Ki-67(139E) | Generated in this study | Figure S7 | 817 |

All constructs are in IRESpuro2 vector backbones.

**Supplemental Table 3. Ki-67 beads charge distribution.**

| Bead | Name | Charge Dephosphorylated | Size (aa) | S | T | Fraction S+T | Phosphorylated S+T | Charge Phosphorylated |
|---|---|---|---|---|---|---|---|---|
| 1 | Nterminal | 3.127 | 135 | 15 | 7 | 0.041 | 6.180 | -9.233 |
| 2 | FASI1 | 2.703 | 120 | 17 | 6 | 0.043 | 6.461 | -10.218 |
| 3 | FASI2 | 4.459 | 120 | 13 | 7 | 0.037 | 5.618 | -6.777 |
| 4 | FASI3 | -0.774 | 120 | 12 | 13 | 0.047 | 7.022 | -14.819 |
| 5 | CP | 24.152 | 186 | 24 | 8 | 0.060 | 8.989 | 6.175 |
| 6 | UpRep1 | 9.147 | 107 | 9 | 7 | 0.030 | 4.494 | 0.158 |
| 7 | UpRep2 | 0.953 | 107 | 15 | 12 | 0.051 | 7.584 | -14.216 |
| 8 | UpRep3 | 3.197 | 107 | 4 | 9 | 0.024 | 3.652 | -4.107 |
| 9 | Rep1 | 2.707 | 121 | 8 | 14 | 0.041 | 6.180 | -9.652 |
| 10 | Rep2 | 4.224 | 122 | 6 | 12 | 0.034 | 5.056 | -5.888 |
| 11 | Rep3 | 0.437 | 122 | 7 | 13 | 0.037 | 5.618 | -10.799 |
| 12 | Rep4 | 4.676 | 121 | 9 | 18 | 0.051 | 7.584 | -10.492 |
| 13 | Rep5 | 6.464 | 121 | 9 | 13 | 0.041 | 6.180 | -5.895 |
| 14 | Rep6 | 5.704 | 122 | 14 | 14 | 0.052 | 7.865 | -10.026 |
| 15 | Rep7 | 3.225 | 122 | 7 | 16 | 0.043 | 6.461 | -9.696 |
| 16 | Rep8 | 4.467 | 122 | 6 | 12 | 0.034 | 5.056 | -5.645 |
| 17 | Rep9 | 0.467 | 122 | 10 | 15 | 0.047 | 7.022 | -13.578 |
| 18 | Rep10 | 6.433 | 118 | 6 | 17 | 0.043 | 6.461 | -6.488 |
| 19 | Rep11 | 4.984 | 121 | 6 | 12 | 0.034 | 5.056 | -5.128 |
| 20 | Rep12 | 1.228 | 122 | 6 | 12 | 0.034 | 5.056 | -8.884 |
| 21 | Rep13 | 5.467 | 122 | 15 | 13 | 0.052 | 7.865 | -10.263 |
| 22 | Rep14 | 6.678 | 120 | 10 | 14 | 0.045 | 6.742 | -6.805 |
| 23 | Rep15 | 0.469 | 119 | 9 | 13 | 0.041 | 6.180 | -11.890 |
| 24 | Rep16 | 5.226 | 110 | 8 | 12 | 0.037 | 5.618 | -6.010 |

Considering 150 phosphorylations, -2 net charge per phosphorylated residue.
